# Environmental complexity modulates information processing and the balance between decision-making systems

**DOI:** 10.1101/2024.03.11.584503

**Authors:** Ugurcan Mugan, Samantha L. Hoffman, A. David Redish

## Abstract

Behavior in naturalistic scenarios occurs in diverse environments. Adaptive strategies rely on multiple neural circuits and competing decision-systems. However, past studies of rodent decision-making have largely measured behavior in simple environments. To fill this gap, we recorded neural ensembles from hippocampus (HC), dorsolateral striatum (DLS), and medial prefrontal cortex (mPFC) while rats foraged for food under changing rules in environments with varying topological complexity. Environmental complexity increased behavioral variability, lengthened HC nonlocal sequences, and modulated action caching. We found contrasting representations between DLS and HC, supporting a competition between decision systems. mPFC activity was indicative of setting this balance, in particular predicting the extent of HC non-local coding. Inactivating mPFC impaired short-term behavioral adaptation and produced long-term deficits in balancing decision systems. Our findings reveal the dynamic nature of decision-making systems and how environmental complexity modulates their engagement with implications for behavior in naturalistic environments.

## Introduction

Naturalistic behaviors often occur in variably complex and uncertain environments, and therefore the ability of animals to adapt their behaviors to changing environments is crucial for their evolutionary success. For example, a busy intersection requires more complex decision-making than an empty racetrack. Similarly, navigating through undergrowth with multiple paths versus grassland with relatively uniform terrain requires different decision processes. Past research has shown that animals (including humans) adapt their behavioral strategies based on experience, environmental complexity, and environmental geometry^1–3^—the balance between decision systems have to adapt to accommodate changing environment and task demands.^4–6^ Furthermore, in rodents, environmental richness has been shown to affect brain structure and overall cognition.^7–9^ Neurophysiologically, adaptive decision-making is mediated by circuit dynamics that are distributed and coordinated across multiple brain areas that represent different components of these complex conditions.^10–19^ Yet, the role that environmental complexity plays in adaptive decision making and the role of environmental demands on the interactions between multiple brain regions is often overlooked and not well understood.^6,20^ To bridge this gap, we simultaneously recorded from multiple, interconnected brain regions on a foraging decision task with changing reward rules across a systematic variation of environment complexity.

Decision-making is thought to arise from multiple, parallel systems in the brain. Deliberation (prospective, planning, model-based decision-making) relies on internally generated models of the environment and expectations of the future.^21–24^ Conversely, the procedural (habit, model-free) system uses learned and cached situation-action associations.^25–27^ Therefore, while deliberative behaviors are flexible and amenable to novel situations (on a familiar environment),^28,29^ procedural behaviors, while computationally efficient, are often inflexible.^11,15,16,30,31^ Previous evolutionary analyses have found that environmental complexity and topology affect the usefulness of these strategies.^3,6^ However, it remains unclear how environmental parameters and geometry influence the neural representations that are used to select between candidate actions, and how and where information is integrated to guide action selection.

Three regions---hippocampus (HC), dorsolateral striatum (DLS), and medial prefrontal cortex (mPFC)--- are hypothesized to make distinct contributions to the decision-making process. HC is hypothesized to encode the state-space and thus enable flexible behaviors^11,15,16,32–35^, while DLS is hypothesized to encode inflexible action chains and thus enable automated behaviors.^27,36–43^ The hypothesis that decision-making arises from at least two competing systems raises questions about how the deliberative (HC-mediated) and procedural (DLS-mediated) systems are balanced to create adaptive behaviors.^12,17,24,44,45^ Dorsomedial prefrontal cortex (mPFC) has been shown to encode various task relevant features, such as the problem space and strategy, and has been put forward as a candidate region responsible for balancing between these parallel decision-systems.^46–53^

Research suggests that HC is a crucial component of the network supporting memory function, cognitive, spatial and non-spatial information.^11,15,16,54–59^ Furthermore, pyramidal cells in HC (‘place cells’) have been shown to be tuned to different environmental parameters such as geometry and space.^15,60–64^ ln addition to spatially-specific firing at the animal’s current position, decoding methods applied to the HC ensemble activity have elucidated time-compressed sequential extra-field firing at decision times — such as during single cycles of theta oscillations — that suggest imagination of future outcomes.^21,65–73^ Thus, HC is a key component of prospective thinking, future planning, and deliberative decision-making. However, it is still an open question as to how environmental geometry changes HC network-level representations, and thus how it influences planning and deliberative systems.

In contrast, dorsolateral striatum (DLS) is hypothesized to encode cached action sequences for procedural learning and decision making.^26,27,30,38,74^ Prior research has shown punctate bursting activity by DLS ensembles that occur at the start and end of action sequences.^36,40,45^ Furthermore, decoding of striatal activity during these bursts has shown that they reflect the current state and past experiences more than future options.^75^ Together these results suggest that DLS is involved in initiating stereotyped behaviors associated with procedural decision-making. However, it is still an open question as to how environmental geometry changes these firing patterns, and thus how it influences procedural responses.

While there have been numerous studies looking at procedural and deliberative decision-making, and their neural instantiations, existing evidence has not addressed the role that task and environmental structure play in modifying the usefulness of procedural or prospective decision-making in differently-complex environments and the balancing between them. Many behavioral tasks commonly used to investigate these questions involve spatial tasks that are very simple (e.g., T-mazes). However, to understand behavior and neurobiology in naturalistic settings, we need to go beyond this simplicity.

To elucidate the arbitration between decision-making processes within complex environments, we developed a dynamic, complex spatial foraging tasks for rats. We used a variant of the left/right/alternation task, a spatial rule-switching task in which rats are required to adjust their behavioral strategies after uncued changes to the reward contingency.^45,46,53,69,76–82^ We quantitively varied the complexity to explicitly evaluate its impact on behavior and neural activity. Using simultaneous silicon probe recordings of HC, mPFC, and DLS ensembles in freely behaving rats, we found that environmental geometry impacted task-relevant neural representations and changed the timing of the transition between decision-making systems. Furthermore, chemogenetic inactivation of dorsal mPFC impaired the ability of the rats to recognize task regularity and stability, and led to both acute and long-term learning deficits, thus further highlighting mPFC’s role in balancing between deliberative and procedural decision-systems.

## Results

### Environmental complexity differentially impacted the deliberative and procedural systems

To study the neural instantiations and representations of the deliberative and procedural decision-making systems and the balancing between them, we used a left/right/alternation (LRA) foraging task (Fig. 1A, Supplementary Fig. 1A, B),^45,46,53,69,76–82^ and collected pairs of simultaneous recordings from dorsal HC CA1, dorsal mPFC, and DLS (Fig. 1B, Supplementary Fig. 1C, D; n_HC_ = 7; n_DLS_ = 4; n_mPFC_ = 7). Using silicon probes, we recorded a total of 4292 single units from HC, 2293 from DLS, and 2609 from mPFC (Fig. 1B, Supplementary Fig. 1C, D, Supplementary Table 1). To study the role of mPFC in selecting the appropriate decision-system, in a separate cohort of rats, we chemogenetically inactivated dorsal mPFC (CaMKlla-hM4Di DREADDs; see Methods) (Fig. 1C, Supplementary Fig. 7A, B; n_DREADDs Virus_ = 8, n_Control Virus_ = 8).

**Figure 1:**
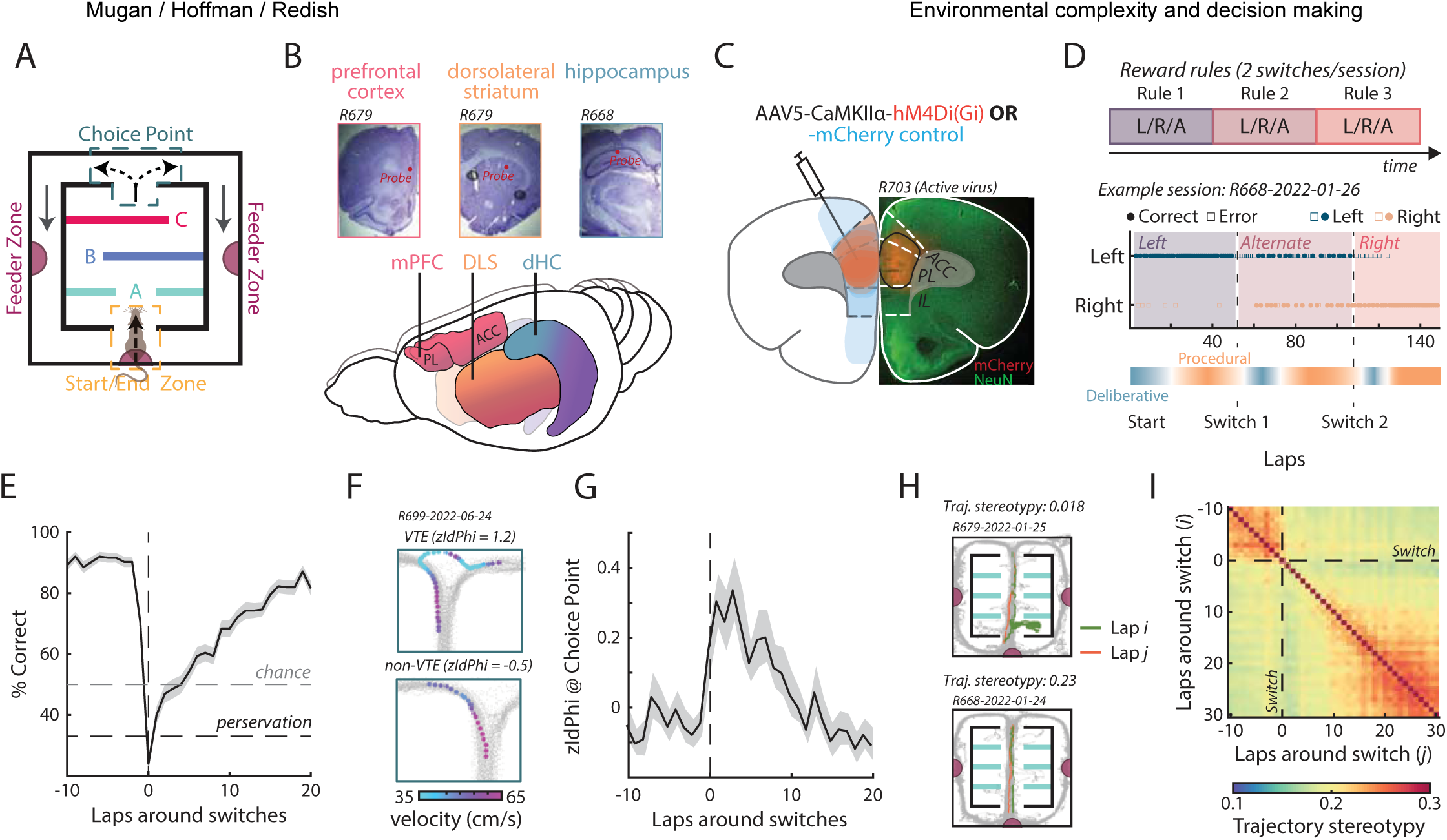
Behavior in left/right/alternation task and environment complexity. **(A)** The left/right/alternation (LRA) foraging task. Three different wall types: open middle (Wall A), open left side (Wall B), and open right side (Wall C) were used to create intermediary decision-points. **(B)** Recording sites from dorsal mPFC, DLS, and dorsal HC. **(C)** Example histology (right) and illustration of viral spread (left) for DREADDs virus animals (red) and control virus animals (blue). **(D)** Example behavioral data from a single session and the hypothesized arbitration between deliberative and procedural decision-making systems. **(E)** Rat performance aligned to rule switches. **(F)** Example vicarious trial and error (VTE) and non-VTE paths. **(G)** z-Scored ldPhi aligned to rule switches. **(H)** Two example paths (orange and green) through the central maze segment showing low (top) and high (bottom) path stereotypy. **(I)** Trajectory stereotypy (reciprocal of Euclidian distance) between pairs of paths through the central maze segment around rule switches.

In the LRA task, rats had to traverse a central maze segment that included three low-cost decision points with dead ends on either side, followed by a high-cost choice point between the left and right return arms (Fig. 1A, Supplementary Fig. 1A).^45,46,53,69,76–82^ There were three rules that determined which return arm would be rewarded as a function of the behavior of the rat. Under the left rule (L), food reward was presented on the left side; under the right rule (R), food reward was presented on the right side; and under the alternation rule (A), food reward was presented on the opposite side of the previous run --- i.e., the rats had to alternate between left and right to receive a food reward. If the rats made the correct decision under the active rule, they received food at the reward arm and again at the start of the maze. Conversely, if the rats made an incorrect decision, they received no food reward at either location. Over the course of a 45 min session, there were two un-cued rule switches per session that occurred roughly at 15 and 30 mins. Each session included all three rules; thus, sessions were drawn from one of six possibilities: LRA, LAR, RLA, RAL, ALR, ARL (Fig. 1D).

In line with previous research,^45,46,53,69,76–82^ rats rapidly adjusted their behavior to the new rule (Fig. 1E, Supplementary Fig. 2A) and performed similarly under each rule with a slight decrease in performance under the alternation condition (Supplementary Fig. 2B, C; mean ± stdv 45.7 ± 12.9 laps/block; mean ± stdv 82.7 ± 10 %correct L & R and 70.3 ± 12 %correct Alt.).

This task is known to probe the interaction between deliberative and procedural decision-making and the neural correlates of their underlying systems (Fig. 1D).^47,53,81,83,84^ Prior work has found that rats display vicarious trial and error (VTE) behaviors around rule switches at high-cost choice points. VTE behaviors entail pausing and orienting back and forth towards the two arm options before making a decision — quantified by ldPhi or (when z-scored) zldPhi (Fig. 1F, G, Supplementary Fig. 2D; see Methods).^24,85,86^ Neural recordings indicate that VTE is often accompanied by HC cell assemblies sweeping forward of the animal serially assessing each option, and thus is believed to reflect a behavioral correlate of deliberation.^21,65,69,77^ In line with previous research, we found that VTE behaviors significantly increased after rule switches (Fig. 1G), which correspond to periods that are hypothesized to be driven under the deliberative system.^46,79,80^

In contrast, prior work has shown that rats develop procedural responses on this task — i.e., behavioral stereotypy — under stable contingencies and after the contingency has been learned.^37,45,75,84^ We quantified behavioral stereotypy by measuring the reciprocal of the Euclidean distance between pairs of path trajectories through the central maze segment on a lap-by-lap basis (Fig. 1H; see Methods).

Notably, rats developed stereotyped path trajectories as they adopted the appropriate behavioral strategy (Fig. 1l). Both the pre-switch and post-switch periods were characterized by an increase in path similarity, with the rule changes disrupting the developed stereotypy.

Prior computational research suggests that environmental structure and complexity should impact the utility of using deliberative vs procedural strategies.^5,6^ Here, we modulated environmental complexity by parametrically changing the central maze segment (central path) (Fig. 2A, Supplementary Fig. 1A). Intuitively, high complexity environments are those that have highly variable navigational path lengths within the central segment. To quantify environmental complexity, we discretized the maze and created a graph of the discretized space. Structural complexity was defined as the total information content across all sets of equivalent trees that could be created from a given maze configuration (see Methods).^87–90^ Walls used to manipulate the interior navigational sequence (A: open middle, B: open left, C: open right) created 27 unique mazes with different levels of complexity (Fig. 2A). These mazes roughly formed three clusters of complexity, which we partitioned into low, mid, and high complexity for subsequent analysis (Fig. 2A).

**Figure 2:**
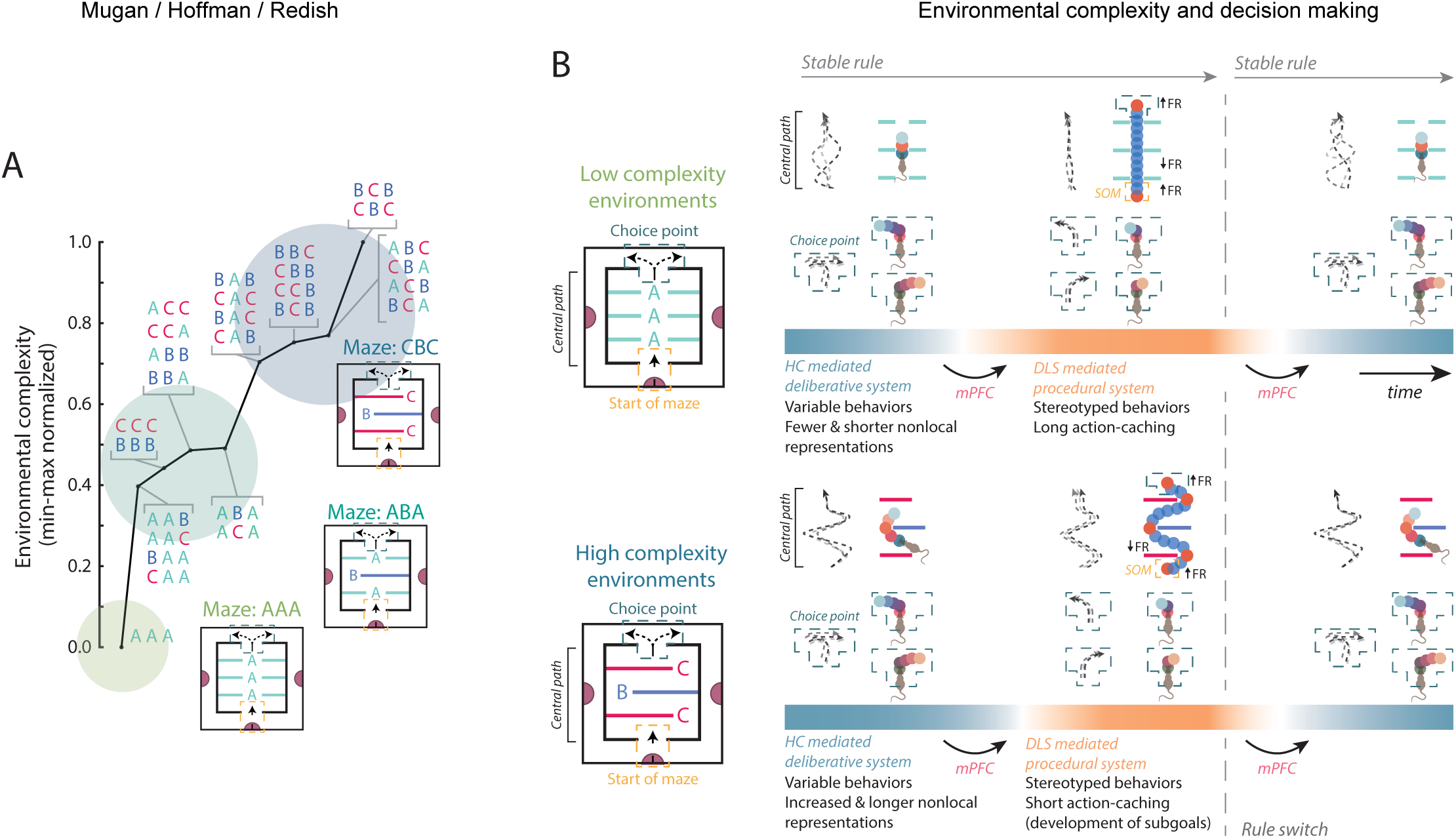
Quantification of environmental complexity. **(A)** Environmental complexity of different maze structures created by rearranging inserts shown in (A). The colored circles indicate the low, mid, and high complexity designations. **(B)** lllustration of hypothesized behavior and neurophysiology in differently-complex environments. For panels E-G, shaded error bars show mean ± SEM. The vertical dotted lines correspond to the last lap of the previous rule block.

Theoretically, if rats are using different decision-making algorithms, they should show predictably different behavioral outcomes, neurophysiological representations, and behavioral-neurophysiological interactions based on the internal structure of the maze. Environments that feature multiple decision-points (low-cost decision-points aligned to walls B and C) are hypothesized to encourage decision-making that relies more on deliberation, thus resulting in more variable behaviors, longer hippocampal nonlocal sequences, and differential engagement of dorsolateral striatum at these low-cost decision-points. In contrast, low complexity environments are hypothesized to encourage decision-making that relies more on habit, thus resulting in the development of stereotyped responses that span the entire central path, with shorter, less sequential hippocampal nonlocal sequences (Fig. 2B). Thus, this environmental control provides us with an experimentally tractable method for incorporating environmental complexity into a behavioral task that is well established for studying the engagement and competition between decision-systems.

To directly assess the role environmental complexity plays in modulating decision systems, we first analyzed the impact of changing complexity on overall task performance. We found that environmental complexity significantly affected performance: rats made more errors (poorer performance) in more complex environments (Supplementary Fig. 2E; Mixed effects ANOVA F(18) = 2.35, p = 0.0033). While there were significant interaction effects between environmental complexity and performance across contingency and rule blocks (Mixed effects ANOVA laps/contingency*complexity F(36) = 2.85, p < 0.001; Mixed effects ANOVA performance/contingency*complexity F(36) = 2.16, p < 0.001) there was no systematic pattern (Supplementary Fig. 2F, G). Looking at the effect of environmental complexity on VTE we found that there was a slight increase in VTE on only high complexity environments (Fig. 3A; Mixed effects ANOVA F(18) = 1.59, p = 0.072, Levene’s test F(2, 141) = 0.79, p = 0.45; no significant difference in the variances across the three complexity groups).

**Figure 3:**
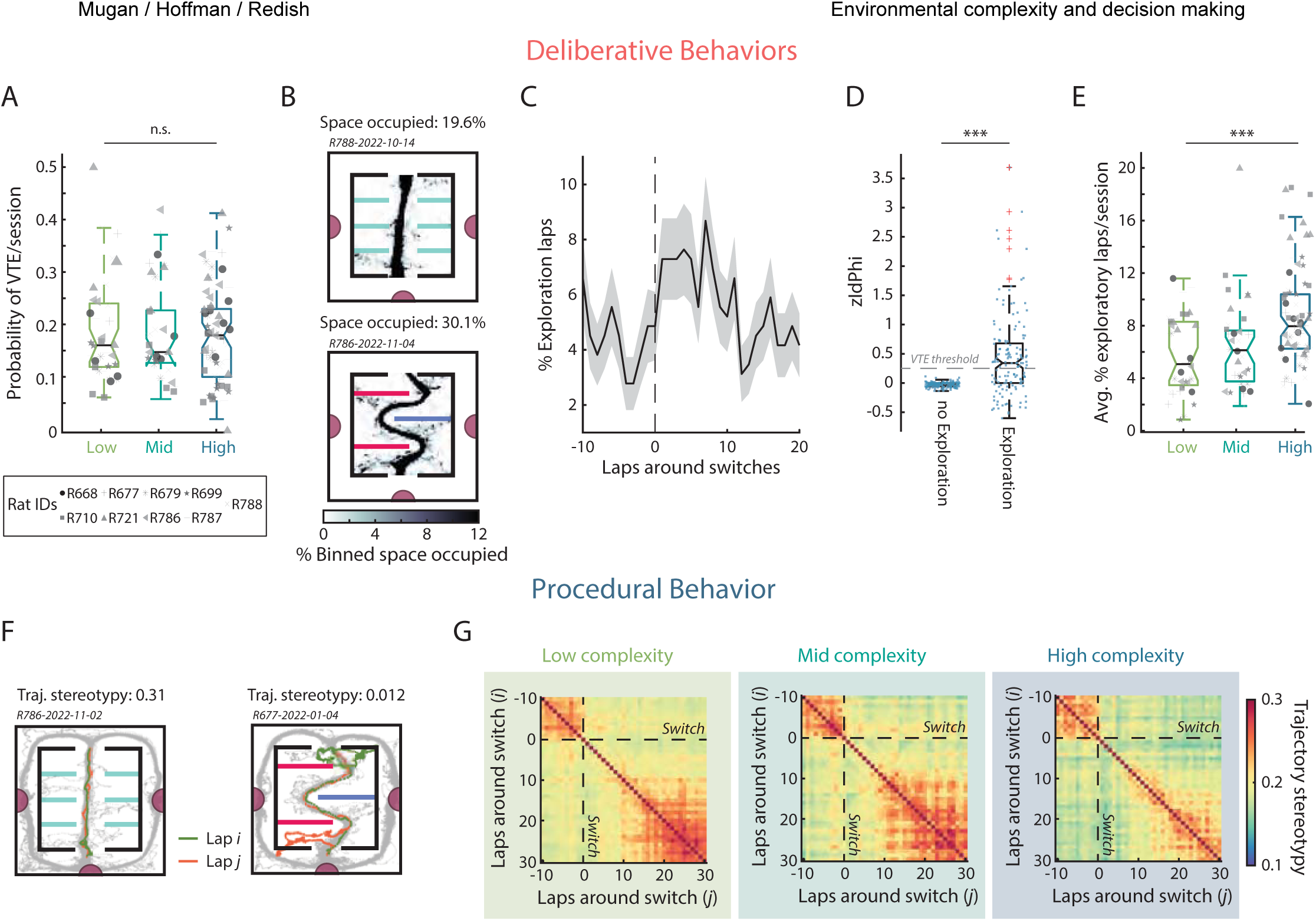
Behavioral elfects of environmental complexity on the LRA task. **(A)** Proportion of VTE laps (zldPhi > 0.25) in low, mid, and high complexity environments. **(B)** Two example sessions showing the occupancy of the central maze segment. *Top*: example of a low exploration session, and *Bottom*: an example of a high exploration session. **(C)** Percentage of exploratory laps aligned to rule switches. Black line indicates mean, and the shaded gray region ± SEM. The vertical dotted line corresponds to the last lap of the previous rule block. **(D)** Distribution zldPhi values at the choice point for non-exploratory and exploratory laps as measured on the central track. Horizontal line denotes the boundary for identifying VTE events (Kruskal-Wallis test H{1) = 46.41, p < 0.001). **(E)** Percentage of exploratory laps per session broken down by maze complexity group. **(F)** Examples of trajectory stereotypy in different environments (as in Fig. 1H). **(G)** Development of stereotypy around rule switches (as in Fig. 1l).

Rats explored the central maze segment (Supplementary Fig. 1A) by occasionally wandering on unrepeated paths to the corners and edges of the dead-end spaces (Fig. 3B). These explorative events increased around rule switches (Fig. 3C, Supplementary Fig. 2H, l) and were correlated with the prominence of reorientation behaviors (Fig. 3D, Supplementary Fig. 2J). These data suggest that exploration also provides a behavioral correlate of deliberation that, unlike VTE, is associated with the central path. Notably, the proportion of exploratory laps increased with environmental complexity (Fig. 3E; Mixed effects ANOVA F(18) = 2.61, p = 0.0011), which is in line with the idea that environmental complexity modulates the competition between deliberative and procedural decision-making.^6^

Furthermore, we found that central path stereotypy — a behavioral marker for procedural decision-making — was also modulated by complexity. Increased complexity resulted in lower behavioral stereotypy and slower development of stereotypy following rule switches (Fig. 3F, G). Concurrently, we found that path stereotypy in the return rails (choice point exit to start of maze entry) decreased (Supplementary Fig. 2K), and side feeder linger times increased after rule switches (Supplementary Fig. 2O), with errors inducing more deviation from stereotyped paths (Supplementary Fig. 2L) and longer feeder wait times (Supplementary Fig. 2P). However, neither metric was modulated by side choice (Supplementary Fig. 2M, Q) or environmental complexity (Supplementary Fig. 2N, R). Therefore, as predicted by previous computational work,^6^ we found that the effects of environmental complexity on behavioral stereotypy was restricted to the central segment of the maze, where, unlike the return rails, there exists variable navigational paths (Fig. 2A).

Taken together these results indicate that modulating the environmental structure has important consequences on deliberative and procedural behaviors.

### Hippocampal sequences were modulated by environmental complexity

Place cells commonly found within the CA1 region of the HC have been shown to provide direct insight into how the brain’s deliberative decision system represents important information.^21,24,29,91–94^ Place fields appeared in both simple and complex environments (Fig. 4A, B), and the number of place cells and their distribution through the maze did not differ across the different environments (Supplementary Fig. 3A, B). As expected, the firing of putative pyramidal cells (89.3% recorded HC units) was largely local throughout the maze (Fig. 4B). However, place fields were significantly smaller in more complex environments, suggesting differences in network-level representations of space in differently-complex environments (Supplementary Fig. 3C; One-way ANOVA F(2) = 9.91, p < 0.001). These differences were not driven by differences in speed through the central path (Supplementary Fig. 3D; Mixed effects ANOVA F(18) = 1.29, p = 0.20). To further investigate how these network level differences may affect nonlocal HC representations, and putative physiological measures of deliberation, we looked at theta timescale (hundreds of millisecond) sequences (Fig. 4C).

**Figure 4:**
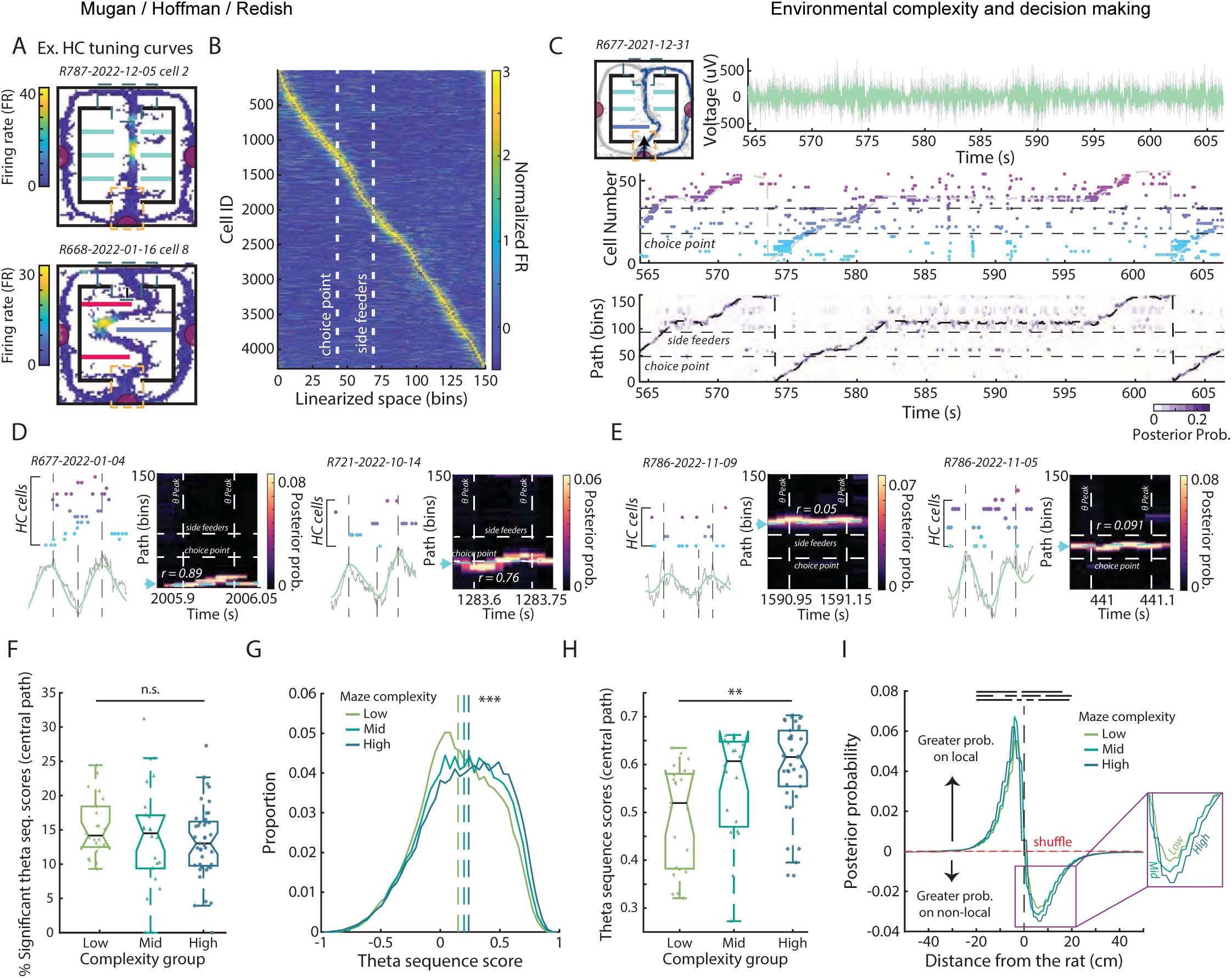
Theta sequences in hippocampus in dilferently-complex environments. **(A)** Two example HC cell tuning curves. **(B)** Spatially binned activity of putative pyramidal cells on the linearized maze (∼3cm bins). **(C)** An example of HC ensemble representation of space. *First row*: The maze and an example trajectory (dark blue) along with raw (gray) and theta frequency filtered LFP (green). *Second row*: spike raster of simultaneously recorded HC cells on an example lap. Cells are ordered and colored by their respective tuning curve centers. Gray line shows the actual trajectory of the rat. *Third row*: Decoded activity from the HC ensemble. Black line shows the actual trajectory of the rat in linearized space. Dashed horizontal lines indicate the high-cost choice point and feeder locations. **(D, E)** Examples of (D) high and (E) low score theta sequences. *Left*: HC cell spiking during a theta cycle ordered and colored by field location. Raw LFP (gray) and theta filtered LFP (green) are shown below with vertical lines indicating the peaks and trough of the theta cycle. *Right*: Bayesian decoding during the theta cycle. Theta sequence scores are indicated on the plots. The arrow and cyan line show the actual position of the rat during the theta time window. **(F)** Average percent of significant theta sequences per session. **(G)** Distribution of theta sequence scores for the different maze complexity groups. Dashed lines indicate medians. **(H)** Session average theta sequence scores along the central path for the different maze complexity groups. **(I)** Differences in posterior probability of the decoding in the 1^st^ half (peak to trough) and 2^nd^ half (trough to peak) of each identified theta cycle (1^st^ half –2^nd^ half) aligned to the actual position of the rat. Positive values indicate more local and negative values indicate more non-local decoding. The red line indicates the shuffle distribution. Black lines indicate significant differences between pairs of environment complexity (top line low vs mid, middle line mid vs high; bottom line low vs high). (Repeated measures ANOVA for complexities across positions; lnteraction: F{34) = 51, p < 0.001. Post-hoc on each position bin via two-sided Wilcoxon rank sum corrected for multiple comparisons;).

Theta oscillations (6-12 Hz) are prominent in CA1 and can contain fast time scale sequential activity of place cells.^21,65,72,95–100^ To find significant theta timescale sequential activity in CA1, theta cycles were identified and limited to those that had 25 cells active. The spike trains within each candidate theta cycle were analyzed using Bayesian decoding^101^ (20ms bins) to relate firing activity to underlying behavior, and the correlation between decoded position and time was measured as a metric of sequential activation (theta sequence score).^102–104^ lntuitively, this metric indicates the ordered nature and clarity of the decoded representations in each theta cycle (see Fig. 4D, E for high and low score examples, respectively). Using this method, we found clear significant theta sequences during the entire navigational period.

While the proportion of significantly sequential theta cycles did not differ between environments (Fig. 4F; Mixed effects ANOVA F(12) = 1.23, p = 0.28 and Supplementary Fig. 3E), we found that the average theta sequence score across all the theta cycles significantly increased with complexity (Fig. 4G; Kruskal-Wallis test H(2) = 765.94, p < 0.001). Moreover, the average theta sequence during traversals along the central path increased with complexity (Fig. 4H; Mixed effects ANOVA F(12) = 2.48, p = 0.01). Notably, similar results were found when taking the slope of the decoded sequences (Supplementary Fig. 3E, F; Kruskal-Wallis test H(2) = 66.67, p < 0.001). Collectively these results suggest that increasing environmental complexity produced more robust nonlocal sequence representations in HC.

ln conjunction with sequential activation of place cells, prior research suggests that theta cycles consist of two components: 1) local representation of the current location during the first-half of theta, and 2) a nonlocal sequential sweep ahead during the second-half of theta.^67^ To quantify the prevalence of nonlocality during theta, we used Bayesian decoding to compute the difference in the posterior probability of the decoded location between the second- and first-half of theta aligned to the actual position of the rat (Fig. 4I). Consistent with past work, we found that the first half of theta contained more local information, and the second contained more nonlocal.^73,76,105,106^ Importantly, environmental structure modulated this balance such that nonlocal decoding during the second half of theta was more prominent in more complex environments (Fig. 4I).

Together, these results suggest that environmental structure played a pronounced role in modulating sequential hippocampal activity during theta cycles. Specifically, theta cycles showed more nonlocal activity with longer look-ahead sequences in more complex environments, which likely required increased cognitive and memory load. This network level modulation of HC activity implies important changes to the deliberative decision-making system based on environmental structure, with more complex environments giving rise to stronger neural representations of future options.

### Theta sequence scores were correlated with deliberative behaviors

Given that we saw changes to CA1 theta sequences with increased environmental complexity, we asked whether there was a coherent relationship between HC nonlocal activity and deliberative behaviors. Briefly, is the prominence of nonlocal HC representations indicative of increased deliberative future thinking, as measured through VTE and exploration?

Consistent with this hypothesis, we found that average theta sequence scores at the choice point were significantly correlated to zldPhi values (associated with VTE events) (Fig. 5A; t-test(80) = -6.8, p < 0.001).

ln line with previous findings, theta sequence scores were higher on VTE laps (Fig. 5B; Kruskal-Wallis H(1) = 7.78, p = 0.0053).^53,76^ Similarly, the average theta sequence scores and the amount of exploration in the central path were also significantly correlated (Fig. 5C; t-test t(84) = -6.9, p < 0.001): theta sequence scores were significantly higher on exploratory laps (Fig 4D; Kruskal-Wallis H(1) = 37.94, p < 0.001). These results replicated when we instead examined sequence slopes (Supplementary Fig. 4A-D).

**Figure 5:**
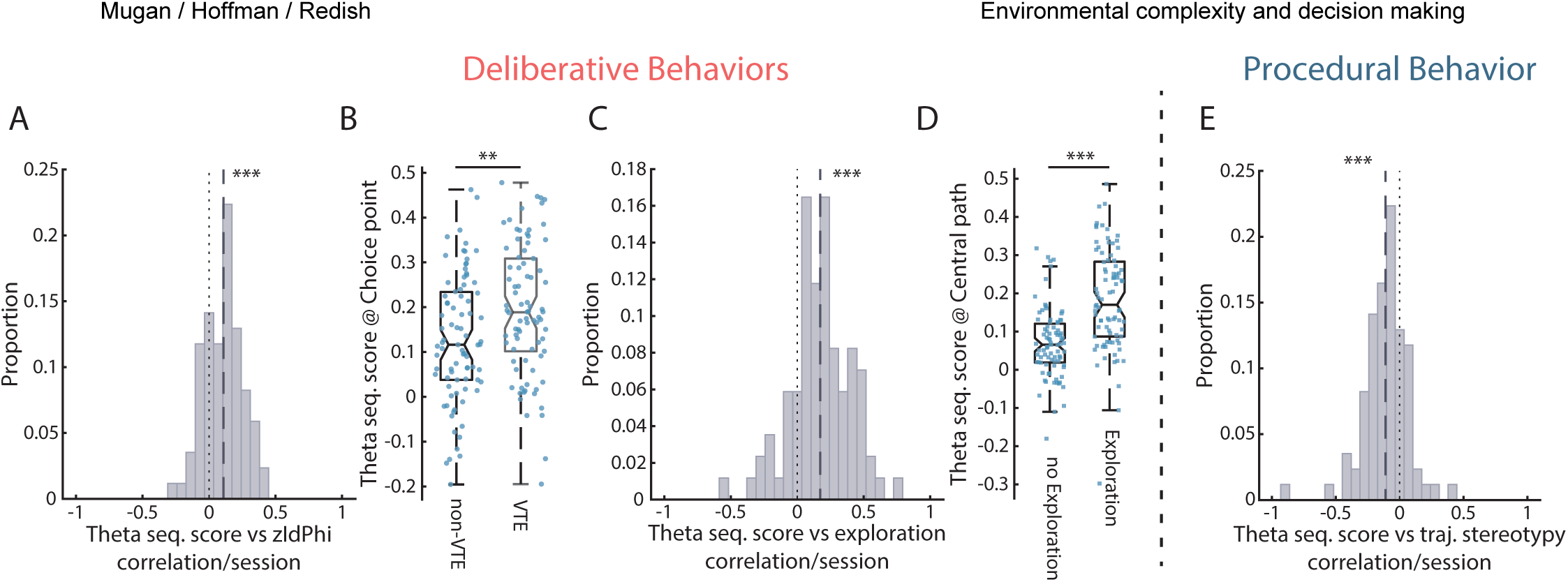
Theta sequence score and its relation to deliberative behaviors. **(A)** Distribution of correlations between zldPhi and mean theta sequence score at the choice point. **(B)** Average theta sequence score at the choice point for laps identified as VTE and non-VTE. **(C)** Distribution of correlations between exploration amount and mean theta sequence score in the central path. Exploration amount was calculated as the average per-lap deviation from the median path in the central maze segment. **(D)** Average theta sequence score through the central path for laps that were identified as exploratory and non-exploratory. **(E)** Distribution of correlations between central path stereotypy and average theta sequence score during central path traversal. (A, C, E) Dashed lines indicate distribution means.

When the HC engages in prospective representations for use in deliberation, we would expect behavior to be less driven by the procedural system. To directly test this, we calculated the average stereotypy of each lap comparing the similarity of its trajectory to prior and subsequent laps (± 5 laps) (see Methods). Notably, trajectory stereotypy and theta sequence scores were inversely correlated (Fig. 5E; t-test t(84) = 5.79, p < 0.001 and Supplementary Fig. 4E), in line with theories that hypothesize a competition between the deliberative and procedural decision-making systems.

These results suggest that CA1 theta sequences accompanied deliberative behaviors, and importantly did not have the same level of sequential structure during procedural behaviors, consistent with hypotheses that theta sequences underlie fast timescale memory recall and deliberation.^73^

### Environmental structure modulated representations of procedural behaviors

After adapting to the new task rule, rats automated their behaviors and showed increased behavioral stereotypy (Fig. 1I, Fig. 3G). This switch to behavioral automation is hypothesized to be driven by dorsolateral striatum (DLS) and is often associated with the development of characteristic firing patterns of DLS cell assemblies, which are hypothesized to reflect the boundaries of well-learned action sequences. These boundaries are believed to indicate the beginning and end of cached action sequences, referred to as “task-bracketing”.^30,36,39–41,107^ To examine the firing pattern of DLS cells, we quantified the average firing rates of putative medium spiny neurons (81.6% of recorded DLS cells) across the linearized maze (mean ± stdv: 2.67 ± 0.03 cm bins and 2.89 ± 0.05 cm bins for the simplest and most complex maze, respectively) per lap for 25 laps aligned to the start of a session and rule switches averaged across the different maze groups (Fig. 6A). Importantly, DLS firing patterns were highly influenced by environmental complexity and structure (Fig. 6A; see Supplementary Fig. 5A, B for lap aligned and average firing rates for each individual maze configuration). In simple environments, which are comparable to other experimental paradigms in which task-bracketing has been studied, such as the classical T-maze, we found action boundaries at the start of the maze and at the choice point (Fig. 6A left panel). Increasing complexity changed DLS activity throughout the navigation sequence. The changes to DLS neuronal activity at discrete positions within the central path across the different mazes were not driven by other motor-related behavioral variables. We found similar residual activity patterns throughout the maze after regressing out either velocity (Supplementary Fig. 6A, B) or ldPhi as a proxy for head turning (Supplementary Fig. 6D, E).

**Figure 6:**
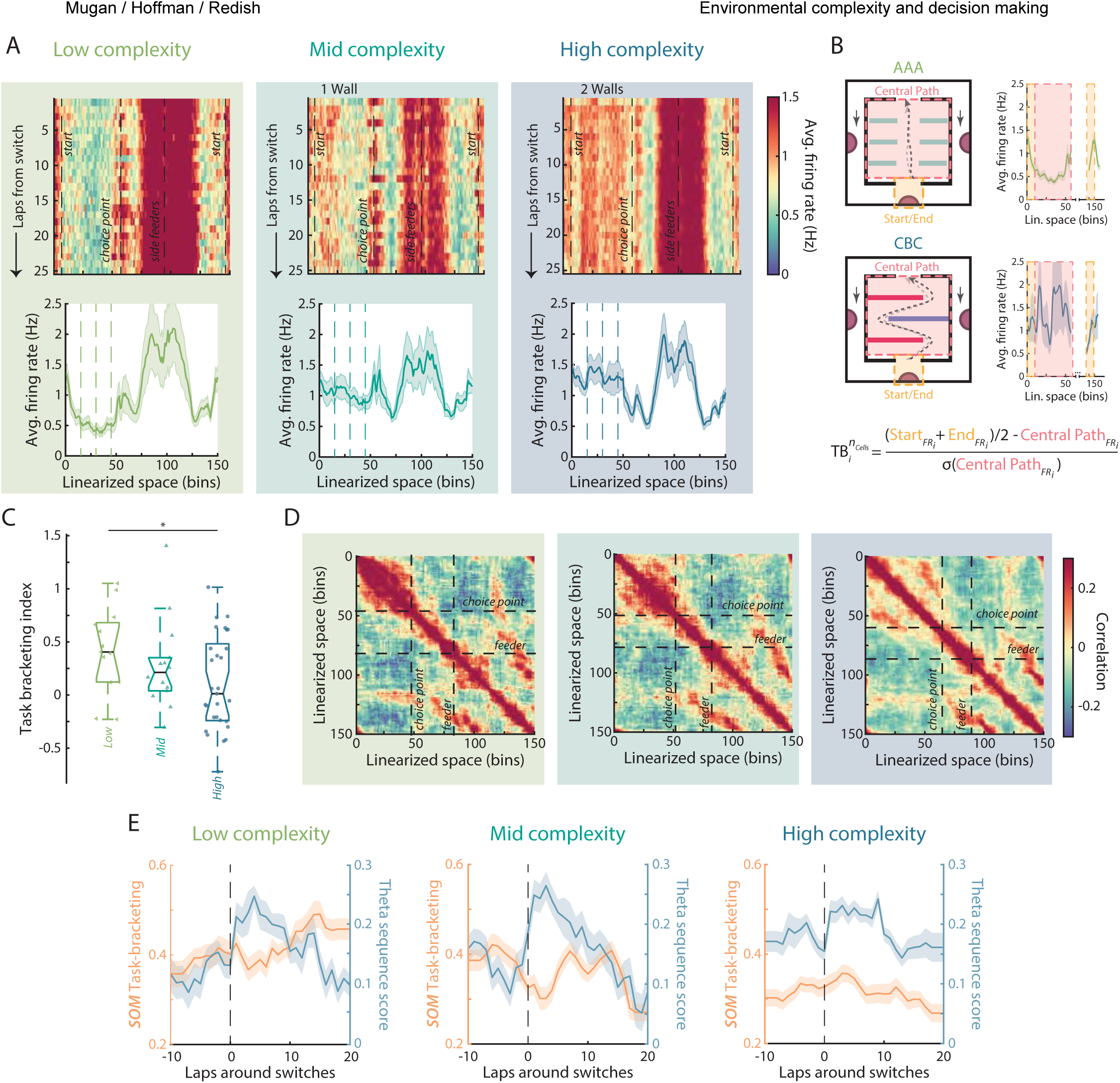
DLS ensemble activity across the linearized maze and the arbitration between deliberative and procedural decision-making. **(A)** *Top row*: Average firing rate of all putative dorsolateral striatum medium spiny neurons (MSNs) over the linearized track (2.67 ± 0.03 cm bins for the simplest maze and 2.89 ± 0.05 cm bins for the most complex maze) averaged across the different complexity groups. Firing rates are plotted as a function of laps aligned to the start of rule blocks. *Bottom row*: Average firing rates averaged across laps and maze group. Solid lines indicate mean, and the shading ± SEM. Dashed lines indicate the internal wall locations projected onto the linearized maze. **(B)** lllustration and equation for the quantification of task-bracketing. **(C)** Average task-bracketing score for each session broken down by maze complexity group. **(D)** Spatial cross correlation of DLS ensemble activity broken down by maze complexity group. The dashed lines indicate the choice point and feeder sites. **(E)** *Blue:* Mean HC theta sequence score across the central path aligned to rule switches. *Orange:* DLS Task-bracketing aligned to rule switches. For both plots, solid lines indicate mean, and the shading ± SEM. The vertical dotted line corresponds to the last lap of the previous rule block

Previous research has quantified task-bracketing — increased firing at the beginning of runs with behavioral automation — as the subtraction of firing rate in the central path from the firing at the start of the maze (task-bracketing index; Fig. 6B).^36,38,39,45^ Positive task-bracketing values suggest the formation of meaningful action boundaries for automation, while negative task-bracketing values indicate that the ensemble firing increases while the action sequence is being implemented and suggest that the action sequence has not been chunked into initiation and termination. Interestingly, we found that task-bracketing index decreased with environmental complexity (Fig. 6C; Mixed effects ANOVA F(8) = 2.31, p = 0.037), as would be expected from slower behavioral automation with complexity.

Given the firing patterns that we observed in mid and high complexity environments, we hypothesized that the structure of the environment might influence how the environment was represented and the boundaries of cached action sequences, as indicated by increased firing at discrete positions in the central path (Fig. 6A, Supplementary Fig. 5A, B). Spatial cross correlation of ensemble activity along a linearized projection of the path elucidates representational similarity between different portions of an environment and provides valuable information of how the environment might be chunked. For example, if ensemble activity is largely local and distinct, one would expect to see high correlation along the diagonal (see Supplementary Fig. 5C for HC example). We found that in simple environments, DLS represented the central path almost as a single entity, where activity along the central path was highly correlated throughout (Fig. 6D left panel, Supplementary Fig. 5D, E; broad block structure), with a representational switch appearing at the choice point (Supplementary Fig. 5D, E). Similar patterning was observed when we instead looked at mPFC activity (Supplementary Fig. 5F). In contrast, with increased environmental complexity, DLS representations became much more local (Fig. 6D averaged across the different maze groups, Supplementary Fig. 5D, E individual maze configurations). Despite increased local representation throughout the central path in both mPFC and DLS we found that wall identity and placement influenced representational similarity. In line with the broad representation of the central path in the AAA maze (Supplementary Fig. 5D-F), we found broad representational similarity throughout the central path in the BBB maze with a choke point occurring between entry into the first wall (Supplementary Fig. 5D-F). In contrast, we found non-uniform representational similarity in more complex mazes when wall identities were not identical. In such cases, environmental structure greatly influenced how the environment was chunked (Supplementary Fig. 5D-F).

We found that HC theta sequences were correlated with deliberative behaviors and inversely correlated with procedural behaviors (Fig. 5E). Concurrently, previous work suggests that task-bracketing is related to the development of procedural behaviors and inversely correlated with VTE behaviors.^5,6,30,36,39–41,107^ Theories hypothesizing a competition between decision systems predict that there should be an inverse relationship between theta sequence scores and task-bracketing, however no one has directly compared these two neural signatures. To compare them, we plotted HC (n_Rats_ = 7) theta sequence scores and DLS (n_Rats_ = 4) task-bracketing aligned to rule changes. We found that there was an increase in theta sequence score and corresponding decrease in task-bracketing aligned to rule switches predominantly in simple environments. While there was a slight dynamic change in DLS task-bracketing in mid complexity environments, there was no corresponding decrease in task bracketing in complex environments, this was because task-bracketing remained low (or never developed) in complex environments (Fig. 6E).

These results highlight the importance of employing varying environmental complexity when looking at the competition between decision-making systems.

### mPFC encoded changes to strategy by engaging the deliberative system around rule switches

The prefrontal cortex is hypothesized to encode the ’problem task space’, goals, subgoals, and strategy.^46–48,108,109^ Accordingly, mPFC has been associated with balancing deliberative and procedural decision-making.^51,110,111^ Previous research has found that mPFC changes its representations around rule switches and behavioral shifts between the two decision systems on the LRA task.^46,47^ One analysis for identifying these changes in mPFC representations has been to correlate lap-by-lap ensemble firing rates in the central path.^46^ In line with previous results, population activity in mPFC was highly consistent when behavior was automated (Fig. 7A left panel), i.e., prior to the switch and after ∼10 laps into a given rule. However, when the rule changed, so too did mPFC ensemble activity.^46,47^ lnterestingly we found similar ensemble dynamics in DLS and HC (Fig. 7A).

**Figure 7:**
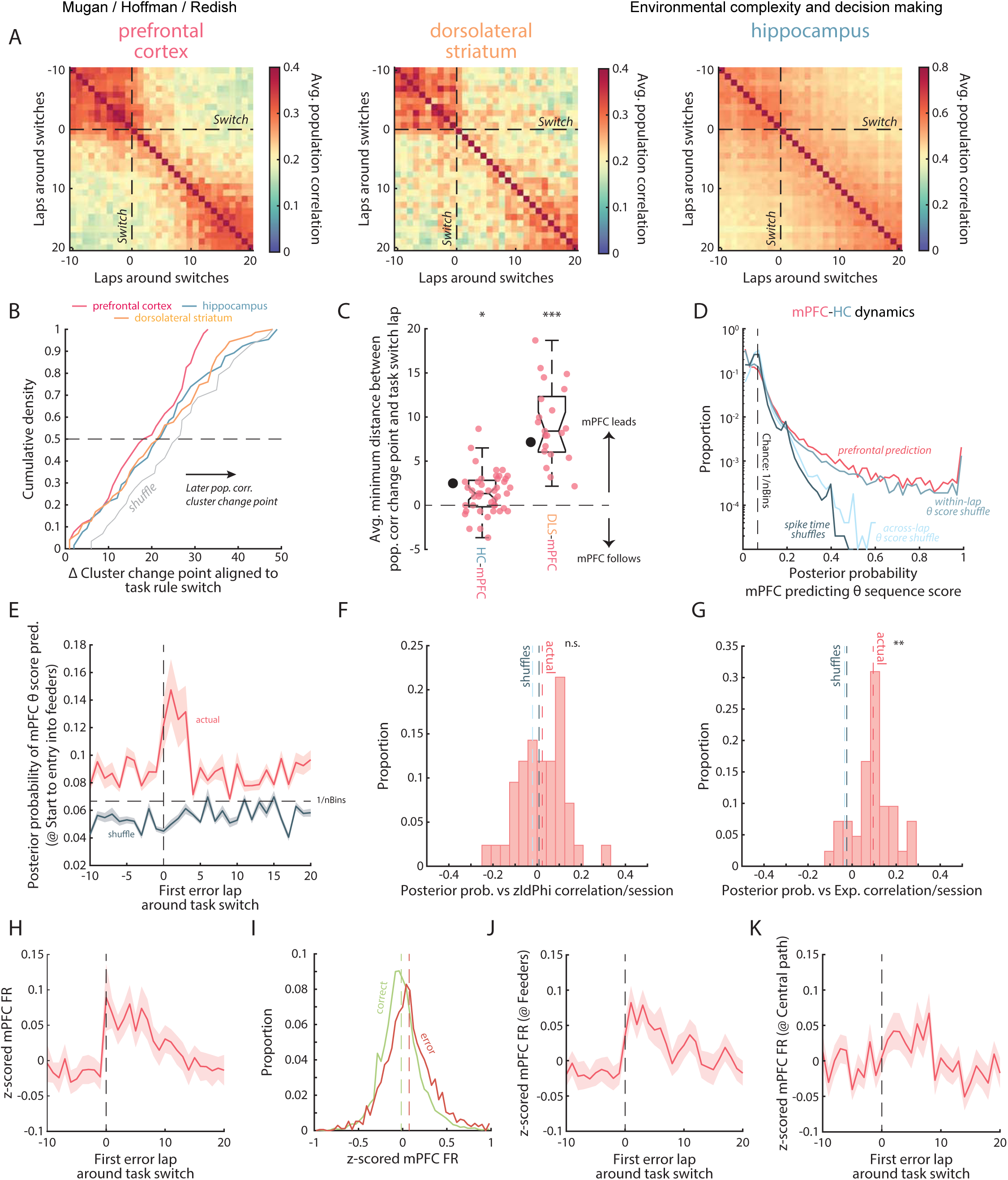
Representational transitions after rule switches in mPFC, HC and DLS and predictions of prospective representations in HC from mPFC activity. **(A)** Lap-by-lap correlation of mPFC, DLS, and HC population activity aligned to rule switches. Correlation matrices are averaged across the rule changes. The vertical and horizontal dashed lines correspond to the last lap of the previous rule block. **(B)** Distribution of physiology-based change points aligned to the task rule switch laps. **(C)** Distribution of the difference in physiology-based change points for simultaneously recorded rats. Black dots indicate mean physiology-based change point lap difference between mPFC and HC and mPFC and DLS in non-simultaneously recorded rats. **(D)** Distribution of the posterior probabilities of mPFC activity predicting binned theta sequence score. The red line indicates actual data. The blue lines indicate the different shuffles. The dashed line indicates uniform probability. **(E)** Posterior probability of mPFC activity predicting binned theta sequence score aligned to the first error lap at or closest to rule switches. Solid lines indicate mean (red: actual, blue: mPFC spike time shuffle), and the shading ± SEM. **(F)** Distribution of correlations between zldPhi and mean posterior probability of mPFC prediction of HC theta sequence score at the choice point. Red dashed line indicates the median of the actual data, light blue line indicates the median of theta sequence score shuffled data, and dark blue line indicates the median of mPFC spike time shuffled data. **(G)** Distribution of correlations between exploration amount (average per-lap deviation from the median path in the central maze segment) and mean posterior probability of mPFC prediction of HC theta sequence score in the central path. Dashed line representation as in (F). **(H)** z-Scored mPFC lap firing rate aligned to the first error lap at or closest to rule switches. Solid line indicates mean, and the shading ± SEM. **(I)** Distribution of z- scored mPFC lap firing rate for correct (green) and error (red) trials. Dashed lines indicate the median. **(J)** z-Scored mPFC firing rate restricted to the side feeder zones aligned to the first error lap at or closest to rule switches. Representation as in (H). **(K)** z-Scored mPFC firing rate restricted the central path (start of maze exit to choice point entry) aligned to the first error lap at or closest to rule switches. Representation as in **(H)**.

If mPFC is responsible for setting strategy, we would expect mPFC to change its firing patterns sooner after a rule switch as compared to either HC or DLS. To identify when ensemble changes occurred, we used a combination of hierarchical clustering to identify reliable ensemble representations and a change point analysis to identify moments that neural activity shifted (see Methods). Across all recordings, representational transitions across all three regions predominantly occurred after rule switches; however, mPFC ensembles changed firing patterns on average earlier than either HC or DLS (Fig. 7B); mPFC representational changes led those in HC and/or DLS, in line with a putative role of mPFC in setting a strategy and relaying this information downstream to HC and DLS.^47^

To directly evaluate the change point timing between regions, we leveraged our simultaneous recordings between pairs of brain areas (mPFC-HC n_Rats_ = 5; mPFC-DLS n_Rats_ = 2). We found that changes to mPFC ensemble firing patterns occurred prior to both HC (mean 1.4 laps; Wilcoxon signed-rank test z = -2.39, p = 0.017) and DLS (mean 9.3 laps; Wilcoxon signed-rank test z = -4.01, p < 0.001) (Fig. 7C).

To study the information transfer between mPFC and HC for strategy setting, we next looked at short timescale prediction of HC state by mPFC activity. To determine whether mPFC activity carried information about what HC was representing, we used a 5-fold cross-validated Bayesian decoding to predict theta sequence scores from mPFC ensemble activity in rats with simultaneous recordings between the two structures (mPFC-HC n_Rats_ = 5; see Methods). Given the positive relationship between theta sequences and deliberative behaviors, mPFC’s prediction of HC nonlocal representations and their sequential structure would elucidate mPFC’s role in switching from procedural to deliberative decision-making.

Critically, we found that mPFC activity predicted the nature of nonlocal representations in HC (Fig. 7D). mPFC activity was more predictive of the HC state compared to: 1) when mPFC representations were destroyed (circularly shifted spike times), and 2) when short-timescale coupling between mPFC and HC state was destroyed, even though internal representations of mPFC were preserved (across-session shuffled HC theta sequence scores). Thus, mPFC representations encoded strategy and could be used to predict HC sequential activity (HC state). However, mPFC did not predict the HC state on a theta-cycle-by-theta-cycle basis. mPFC activity was not more predictive than shuffled theta sequence scores when that shuffling was restricted to a lap (breaking short-timescale relationship between mPFC and HC state while preserving longer timescale relationships) (Fig. 7D).

We found that there was a significant coupling between mPFC activity and HC state, but it was over longer behaviorally relevant timescales (Fig. 7D), suggesting a role in strategy setting on any given lap.^53^ ln looking at the dynamics of this coupling we found that mPFC activity was more predictive of HC state following rule switches (Fig. 7E). Notably, while there was not a significant relationship between prediction posterior probability restricted to the choice point and zldPhi values (associated with VTE events) (Fig. 7F; t-test(41) = -1.18, p = 0.25), we found a significant correlation between mPFC prediction power and exploration amount in the central path (Fig. 7G; t-test(41) = -2.74, p = 0.009), which suggests that these two deliberative behaviors (VTE and exploration) may be associated with different neural processes.

Previous research suggests the existence of reward-related modulation of dorsal mPFC (anterior cingulate cortex) activity.^112–114^ ln line with that research, we found that mPFC activity increased following rule switches (Fig. 7H) and had an overall increase in firing rate following errors (Fig. 7I). Specifically, the increase in firing rate at the feeders (Fig. 7J), in which error signaling would be immediate (omission of expected reward), preceded the modulation of mPFC activity in the central path (Fig. 7K).

These results suggest that mPFC activity and mPFC-HC information flow was modulated by task dynamics, with increased mPFC engagement following rule switches during the hypothesized deliberative periods.

### Inactivating mPFC reduced deliberative behaviors and delayed behavioral adaptations to rule switches

While we found a relationship between mPFC population activity and initiation of strategies, it remained unclear to what extent mPFC causally influenced the balance between deliberative and procedural strategies. To test this question, we used inhibitory DREADDs (designer receptor activated by designer drugs)^115^ in a separate cohort of rats (AAV5-CaMKlla-hM4Di(Gi)-mCherry; n_Rats_=8, n_Sessions_=252; DREADDs) compared to an across-subject viral control (AAV5-CaMKlla-mCherry; n_Rats_=8, n_Sessions_=256; Control) to investigate the behavioral effects of dorsal mPFC inactivation (DCZ^116^ condition) (Fig. 8A, Supplementary Fig. 7A, B; see Methods). To investigate the long-term effects of mPFC inactivation and 2^nd^ day compensatory changes after either mPFC inactivation (hereafter referred to as DREADDs: DCZ) or within-subject control (hereafter referred to as DREADDs: VEH) conditions, we also included a post-manipulation saline repeat of the task (post DCZ or post VEH), which provided us with a washout day (Supplementary Fig. 7C).

**Figure 8:**
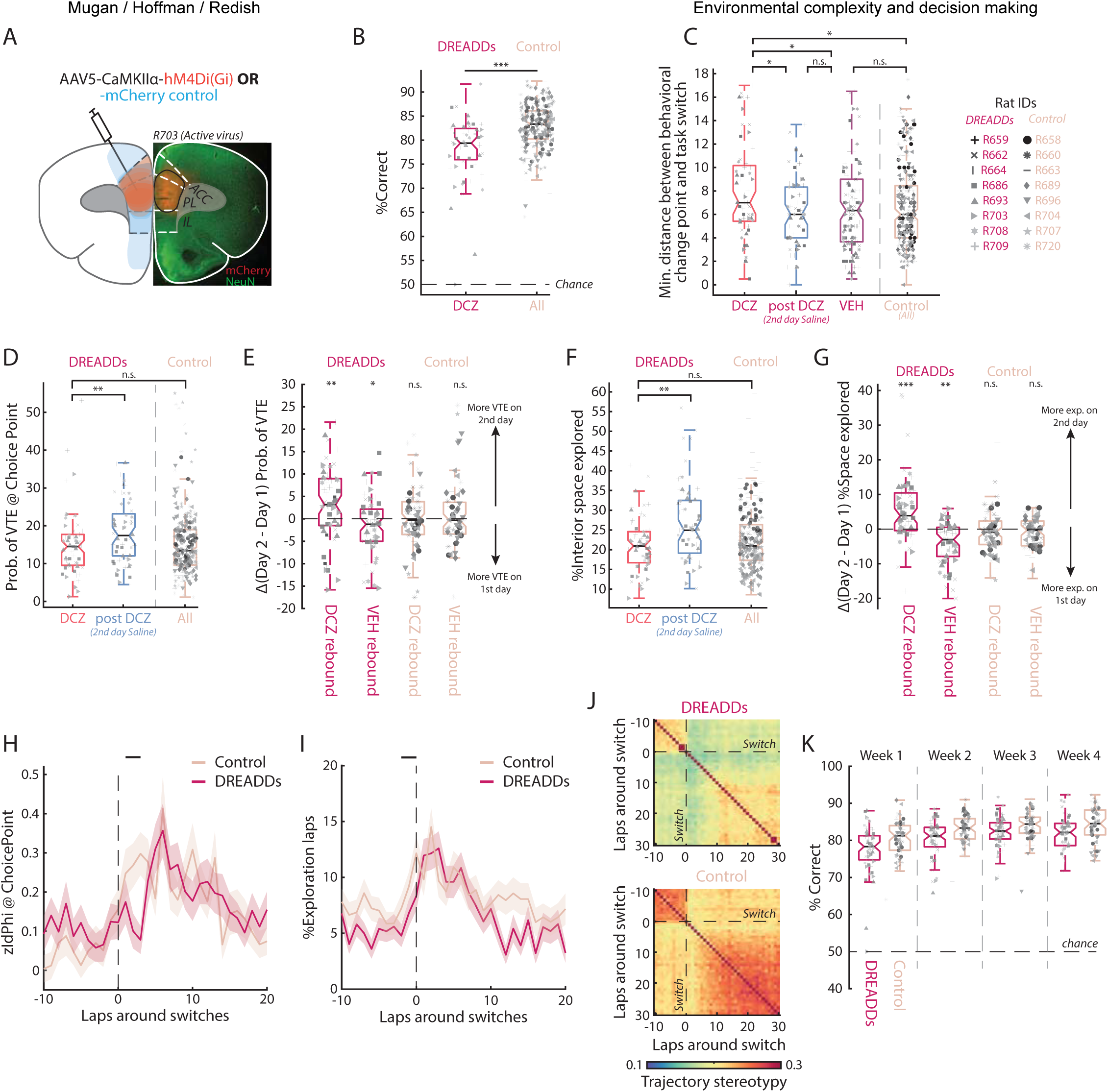
Behavioral elfects of mPFC inactivation with DREADDs. **(A)** Example histology (right) and illustration of viral spread (left) for DREADDs virus animals (red) and control virus animals (blue). **(B)** Distribution of rat performance for mPFC inactivated rats (DREADDs: DCZ dark pink) and control virus animals (light pink). **(C)** Distribution of minimum distance between detected behavioral change points and task rule switch laps for DREADDs virus rats under DCZ, post DCZ (2^nd^ day saline repeat of the task), and VEH, as well as all control virus animals. **(D)** Distribution of proportion of VTE events per session for prefrontal inactivated animals (DREADDs: DCZ, red), for the day after prefrontal inactivation (DREADDs: post DCZ, blue), and for control virus animals (Control, light pink). **(E)** Difference in the proportion of VTE events between 1^st^ day manipulation and 2^nd^ day rebound (Day 2 – Day 1) for DREADDs (dark pink) and control (light pink) virus animals with respect to drug condition. There was a significant virus and DCZ interaction: ANOVA virus*DCZ F{1) = 3.86, p = 0.05. **(F)** Per session distribution of the percentage occupied space in the central segment of the maze for prefrontal inactivated animals (DREADDs: DCZ, red), for the day after prefrontal inactivation (DREADDs: post DCZ, blue), and for control virus animals (Control, light pink). **(G)** Difference in the percentage of the central segment of the maze occupied between 1^st^ day manipulation and 2^nd^ day rebound (Day 2 – Day 1) for DREADDs (dark pink) and control (light pink) virus animals with respect to drug condition. There was a significant virus and DCZ interaction: ANOVA virus*DCZ F{1) = 25.41, p < 0.001. **(H)** zldPhi aligned to rule switches for DREADDs and control virus animals. The horizontal dashed line indicates the threshold for which events were identified as binary VTE and non-VTE events. The vertical dotted line corresponds to the last lap of the previous rule block. The black line above indicates laps in which there was a significant difference between Active and Control groups. Data are shown as mean ± SEM. **(I)** Proportion of exploratory laps aligned to rule switches for DREADDs and control animals. Same display as (H). **(J)** Trajectory stereotypy (reciprocal of Euclidian distance) between pairs of paths through the central maze segment around rule switches for DREADDs and control virus animals. **(K)** Rat performance across the experimental weeks for DREADDs and control virus animals. Each ’week’ is an 8-day sequence (see Supplementary Fig. 8C).

lnterestingly, we found that mPFC inactivation resulted in overall worse performance when compared to the across-subject control virus rats, pointing towards a fundamental involvement of mPFC functioning in proper execution of the task (Fig. 8B, Supplementary Fig. 7D, E; One-way ANOVA F(1) = 22.95, p = 0.0001, n_Sessions-DCZ_ = 64). Critically, the central component of the task is the rats’ ability to adapt their decision-making following a change to the task rule. To quantify how mPFC manipulation changed the speed of behavioral adaptation, we computed the behavioral change point (see Methods). mPFC inactivation (DREADDs: DCZ) led to slower behavioral adaptation to rule switches when compared to post-manipulation saline and VEH days, as well as control virus animals (Fig. 8C; One-way ANOVA F(3) = 3.51, p = 0.015; HSD Post-hoc DCZ vs all others p < 0.05, all other comparisons p > 0.05; n_Sessions-DCZ_ = 64, n_Sessions-postDCZ_ = 64, n_Sessions-VEH_ = 124).

Given evidence that mPFC activity may be influencing deliberative strategies through HC engagement, we investigated the effects of mPFC inactivation on VTE and exploration, behaviors reflective of deliberation and putative HC engagement. Surprisingly, we found that the proportion of VTE events under inactivation (DREADDs: DCZ) was not different from the proportion of VTE events observed in mPFC-intact animals (Control) (Fig. 8D; Mixed effects ANOVA F(8) = 2.58, p = 0.0094; HSD Post-hoc DCZ vs post DCZ p = 0.0038 and DCZ vs Control p = 0.96). Similarly, the amount of space explored after mPFC inactivation was comparable to control virus animals (Fig. 8F; Mixed effects ANOVA F(8) = 6.25, p < 0.001; HSD Post-hoc DCZ vs post DCZ p < 0.001 and DCZ vs control p = 0.47). Thus, while mPFC inactivation impaired the rats’ ability to adapt to new task rules, we paradoxically found no evidence for acute deficits in rats’ ability to employ deliberative behaviors. One possibility is that while rats engaged in deliberative behaviors, inactivation of mPFC impaired their ability to use the information that they gained about the task and rules to perform efficiently.

Building on this hypothesis that mPFC serves a critical role in the use of gained information, we turned to a unique component of our experimental design. By repeating each behavioral condition on a second day in a saline washout condition between manipulations (DCZ rebound n_Sessions-Active_ = 64 and n_Sessions-Control_ = 65, or VEH rebound n_Sessions-Active_ = 62 and n_Sessions-Control_ = 63), we were able to directly evaluate prolonged short-term impacts of mPFC inactivation on task performance and compensatory behavior rebounds following acute mPFC inactivation. Interestingly, the effects of mPFC manipulation were much more pronounced when we considered behavioral differences between these rebound effects. There was a significant compensatory increase in VTE behaviors following mPFC inactivation on the previous day (Fig. 8E DREADDs: DCZ rebound; Wilcoxon signed-rank test z = 2.53, p = 0.0043); but a significant decrease in VTE if mPFC functioning was not manipulated (Fig. 8E DREADDs: VEH rebound; Wilcoxon signed-rank test z = -1.65, p = 0.049 and Supplementary Fig. 7F). Similarly, exploration increased on the second day following mPFC inactivation (Fig. 8G DREADDs: DCZ rebound; Wilcoxon signed-rank test z = 3.63, p < 0.001), but decreased on the second day if mPFC functioning was not manipulated (Fig. 8G DREADDs: VEH rebound; Wilcoxon signed-rank test z = -2.41, p = 0.0079 and Supplementary Fig. 7G). Neither of these compensatory effects after mPFC manipulation were seen in control animals (Fig. 8E; Control: DCZ rebound Wilcoxon signed-rank test z = -0.13, p = 0.9, Control: VEH rebound Wilcoxon signed-rank test z = -0.38, p = 0.7 and Fig. 8G; Control: VEH rebound; Wilcoxon signed-rank test z = -1.63, p = 0.1, VEH condition: Wilcoxon signed-rank test z = -0.18, p = 0.89).

Given the impact that our mPFC manipulation had in eliciting compensatory behavioral rebounds following inactivation, we hypothesized that our manipulation may be causing long-term deficits in strategy setting and learning. To analyze the potential effects mPFC manipulation may have had on deliberative strategies, we compared the changes to VTE and exploration around rule switches in DREADDs (active virus) and control virus animals by pooling across all drug and rebound conditions. In line with our previous findings, we found that both the prominence of reorientation behaviors (zldPhi) and the proportion of exploratory laps increased around rule switches in both DREADDs and control virus animals. Importantly, this increase happened earlier in control animals, especially when looking at zldPhi, suggesting that control animals switched decision-making strategies faster (Fig. 8H; Repeated measures ANOVA around switch interaction F(5) = 2.61, p = 0.023 and Fig. 8I; Repeated measures ANOVA around switch interaction F(6) = 2.12, p = 0.048, Post-hoc on each position bin via two-sided Wilcoxon rank sum corrected for multiple comparisons). While we found significance, the increase in exploration appeared just before rule switches and was very small, suggesting it may be due to noise (Fig. 8I).

To determine if mPFC inactivation affected rats’ ability to employ procedural strategies, we compared the development of behavioral stereotypy around rule switches in DREADDs and control virus animals. We found that DREADDs animals developed behavioral stereotypy slower and showed overall less stereotypy compared to control virus animals (Fig. 8J, Supplementary Fig. 8A), even when their exploration level was the same as control animals (Fig. 8F, Supplementary Fig. 8A DCZ) likely due to subtle differences between paths through the central track. This is in line with previous findings that suggest dorsal mPFC disruption causes long-term deficits in habit formation.^117^

Finally, we evaluated the long-term progression of task performance over the course of our multi-week experiment. Notably, we found that DREADDs animals performed worse throughout the entire experimental sequence (Fig. 8K; One-way ANOVA virus F(1) = 9.9, p = 0.071). In considering how rats’ behavioral strategies may have evolved over the course of the experimental timeline, we hypothesized that as rats became more familiar with the general task structure (rule changes and adaptation), deliberation became less critical. In line with this hypothesis, we found that control rats exhibited progressively less exploratory behavior over the course of the four-week experiment (Supplementary Fig. 8B; Mixed effects ANOVA week F(6) = 3.96, p = 0.0032), though VTE remained stable (Supplementary Fig. 8C; Mixed effects ANOVA week F(6) = 0.7, p = 0.65). In contrast, DREADDs rats did not exhibit this observed decrease in exploration or VTE.

Collectively, these data suggest that mPFC inactivation impaired long-term behavioral adaptation and recognition of task regularities. Furthermore, it impaired effective engagement and balancing between deliberative and procedural decision-making systems. These deficits resulted in both short-term and long-term performance decreases and highlight the importance of mPFC in this system.

## Discussion

Prior research into decision-making suggests that adaptive decision-making necessitates the interaction between multiple competing decision-making systems, two of which are deliberative (or prospective) and procedural (or habit) systems. Current theories suggest that these two systems depend on different neural circuits, are optimized for different situations, and are reflected in behavioral differences during decision-making.^10–12,15,16,118,119^ However, the environment in which these decision-making processes are taking place is often overlooked. In the present study, we examined how behavior, neural representations, and the interactions between the key brain regions of hippocampus (HC), dorsolateral striatum (DLS), and medial prefrontal cortex (mPFC) changed in uncertain and differently-complex spatial environments. The left/right/alternation foraging task with changing internal barriers, coupled with simultaneous recordings across dorsal HC, DLS, and dorsal mPFC, allowed us to directly and quantifiably study the effects of environmental complexity on neural representations and decision-making strategies.

Past research has emphasized the competition between deliberative and procedural control of behavior and has suggested that they arise based on the animal’s experience and the consistency of the environment.^13,16,25,74,120^ Deliberative control is favored when the optimal actions remain variable.^11,15,16,24,121–124^ In contrast, procedural control is favored when the optimal behavior is relatively consistent across time.^11,15,16,36,107,118,125^ In line with this idea, decision-making strategy in our task changed from initially deliberative on entry into the environment to procedural as the rats settled into a behavioral flow, but then transitioned back to deliberative around rule switches before becoming automatic again under the procedural system after adaptation to the new rule. Importantly, environmental structure affected this process, slowing behavioral adaptation in complex environments (Fig. 3A, E, G).

### Competition between deliberative and procedural decision-making in differently-complex environments

The LRA task engages the brain regions hypothesized to play prominent and distinct roles in deliberative and procedural decision-making (HC, DLS, and mPFC) at distinct moments in the task that correspond to hypothesized periods of deliberative and procedural decision-making (Fig. 1).^47,53,81,83,84^

The role of HC is often thought of as providing an adequate model of the environment (or cognitive map) to support future thinking of potential action sequences and their outcomes to aid deliberative decision-making.^11,16,21,24,29,66,69,91,126,127^ We found that time-compressed HC nonlocal sequences predominantly occurred during deliberative behaviors such as VTE (Fig. 5A, B). Importantly, we observed another deliberative behavior, which we identified as exploration, that was also affected by maze complexity and was correlated with the prominence of HC nonlocal sequences, significantly more than VTE, thus further buttressing the role of HC in model-building (Fig. 3B-D, Fig. 5A-D, Supplementary Fig. 4A-D).

Furthermore, complex environments featured more theta-timescale nonlocal activity that were longer and more prominent (Fig. 4). Together these data suggest that prolonged deliberation in complex environments may indicate increased cognitive/memory load due to the need to create fine-scale schema of the environment to learn the consistencies therein.

Prior research has shown that there is a parallel between deliberative decision-making and curiosity, which is often associated with the intrinsic motivation and natural tendency to proactively explore the environment to gather information about its structure.^128^ Thus, this structural knowledge can be flexibly used when planning actions in new environments or when reward locations change.^11,15,28,29,71,93,123,129^ Here, the complexity of the environment, which quantifies the diversity and length of available paths within the central segment of the maze, may be changing the utility of exploration (Fig. 3E, Supplementary Fig. 2H-J). In low complexity environments the ability of the animal to visually observe the high-cost choice point at the end of the central path may be enough for it to infer the structure of the environment, decreasing the overall usefulness of physical exploration. Another possibility is that the diversity of the number of paths increases the perceived state-space of the problem thus favoring increased physical exploration that implicitly organizes the mental search and planning occurring over the state-space.

The difference in the strength of relationship between mPFC prediction of HC state during VTE events and exploration suggests that these two behaviors associated with hypothesized deliberative periods likely reflect different neural processes. Exploration, following initial task entry and rule changes, can be associated with either curiosity or induced novelty. Vicarious trial and error behaviors more frequently occur after a model of the environment has been established, but while there is still uncertainty about the best actions to take to maximize reward outcome.^24^ Exploration may be facilitating the encoding of a new model of the environment, which is then retrieved during VTE events. Unlike exploration, VTE instead could be HC broadcasting information out to connected structures about the established environmental model for valuation to occur in the upstream regions. This would be in line with our finding that shows that while there is no significant relationship between zldPhi (associated with VTE events), there is a significant relationship between exploration amount and mPFC posterior probability (Fig. 7F, G).

Procedural decision-making is often thought to be encoded by DLS bursts aligned to meaningful boundaries of well-learned action sequences.^27,30,36,37^ Consistent with previous research^45,75,107^, we found increased firing at the beginning and end of action sequences in simple mazes (Fig. 6A, C, Supplementary Fig. 5A, B). Task-bracketing and DLS representations of cached action-sequences have primarily been studied in simple environments, such as T-mazes. In more complex environments, DLS ensemble firings were modulated by the internal structure of the maze. Furthermore, DLS representations of space changed with environmental complexity, from highly similar throughout the navigation sequence in simple environments, to local and distinct in complex environments (Fig. 6C).

The dorsolateral striatum receives most of its excitatory inputs from sensorimotor cortex and thalamus, and can influence movements by modulating, for example, motor cortical activity.^130–132^, This raises questions as to cognitive vs motor components in DLS firing patterns. While it is possible that the increased firing and shared representations in the central path in more complex environments may be a byproduct of distinct motor actions, these observations are also consistent with theories that suggest these bursts are indicative of action selection and initiation through the integration of sensorimotor, cognitive, and motivational information.

Our findings suggest that the potential balance between the deliberative and procedural decision-making systems may not be as distinct in environments that require significant cognitive/memory load, and that striatum may not be encoding classical task-bracketing representations in complex environments aligned to task start and end points. The hypothesis that environmental complexity increases memory load is in line with computational research that suggests that planning and deliberation grow quadratically with the state-space.^3^ Task-bracketing in complex environments may not be the right metric to understand how DLS may be representing the boundaries of cached action-sequences.

### The role of dorsal prefrontal cortex in balancing between decision systems

The hypothesis that behavior arises from an interaction between multiple algorithms leaves open the critical question about how disagreements between decision-making systems are resolved. Current theories suggest that these are resolved through information processing in the medial prefrontal cortex.^46–52,133,134^ Consistent with this hypothesis, dorsal mPFC ensemble firing patterns changed following rule switches, and preceded representational changes in both HC and DLS (Fig. 7A-C). At the level of single HC theta cycles, mPFC activity predicted the prominence of HC nonlocal sequences (Fig. 7D). The degree of mPFC predictiveness of HC state and overall mPFC activity was dynamically modulated by task and behavior (Fig. 7E-G). Consistent with the idea that mPFC and HC coordinate to produce adaptive behaviors, prior research has shown that they exhibit oscillatory synchronization during decision-making tasks, which is interpreted as information transfer and integration between mPFC and HC.^22,135–138^ Modulation of mPFC firing rates (Fig. 7H-K) and mPFC leading both HC and DLS in changing its activity patterns following a rule switch, suggests that mPFC may be integrating contextual information to balance between deliberative and procedural decision-making, by updating the current task rule and inhibiting prepotent responses to enable flexible behaviors.

To further examine the role of dorsal mPFC on behavioral adaptation and flexibility, we chemogenetically inactivated dorsal mPFC. We found that mPFC inactivation led to acute performance deficits and slower adaptation to rule changes. This is in line with previous research that has found disruptions in working memory and perseverative errors with dorsal mPFC lesions.^139–141^ Consistent with prior research that has shown that neural activity in anterior cingulate cortex shifts between exploration and choice repetition,^142,143^ we found that dorsal mPFC inactivation caused perseveration of left or right choices after rule changes leading to maladaptive behaviors (Fig. 8C). Surprisingly, mPFC inactivation did not decrease the prevalence of deliberative behaviors when compared to control animals but resulted in long-term changes to strategy setting, suggesting inadequate learning from deliberative behaviors. Importantly, we saw compensatory mechanisms that increased deliberative behaviors following mPFC inactivation (Fig. 8E, G). This suggests that mPFC manipulations may be having long-term strategy effects, where continuous inactivation may be leading to global impairments in building context associations that are necessary for connecting a situation with a decision strategy that is appropriate for the degree of behavioral flexibility perceived to be required.

## Acknowledgements

We thank G. Cannan, T. Lawrence, J. Williams for help with experiments, as well as C. Boldt, K. Seeland, and A. Sheehan for technical assistance. We thank E. Krook-Magnusson, C. J. Walters, and members of the Redish lab for comments on earlier drafts and useful discussions. This work was supported by NlH grants R01-MH112688 and R01-MH080318.

## STAR Methods

### Resource Availability

#### Lead contact

Further information and requests for resources should be directed and will be fulfilled by the lead contact, A David Redish.

#### Materials and availability

This study did not generate new unique reagents.

#### Data and code availability

Data and code required to reproduce the findings reported in this paper will be made available on OSF.

### Experimental model and subject details

#### Subjects

Eight adult Fisher-Brown Norway F1 hybrid rats (FBNF-1) (5M, 3F) and one adult Brown Norway (BN) female were used in the recording study (n = 9 rats). FBNF-1 rats (1^st^ generation hybrid of F344 female and BN male) were bred in-house, and the BN rat was purchased from Envigo. All rats were aged between 6-11 months old during experiments. While two strains of rats were used in the reported study, we found no evidence for differences in behavior or physiology between FBNF-1 and BN rats, so data were combined.

Ten adult BN rats (5M, 5F) and six FBNF-1 (3M, 3F) were used in the manipulation (DREADDs) study (n = 16 rats). All rats were aged between 7-15 months old during experiments. While two strains of rats were used in the reported study, we found no evidence for differences response to manipulations between FBNF-1 and BN rats, so data were combined.

Rats were single housed in a temperature-controlled colony room with a 12 hr light/dark cycle (lights on at 8:00 AM). Throughout the experiments the rats were food restricted and had to maintain a weight of at least 80% of their free feeding weight. Rats were given full access to food one-week prior to probe implantation or DREADDs (designer receptors activated by designer drugs) viral infusion. Surgeries were conducted when the rats were at least at 100% of their prior free feeding weight. All training and experimental sessions were conducted at around the same time and during the rats’ light phase. All procedures were approved by the University of Minnesota lnstitutional Animal Care and Use Committee (lACUC) and were in accordance with the NlH guidelines.

### Method details

#### Surgical procedures

Of the 25 rats used in the study, 9 rats were used for *in vivo* electrophysiology recordings, and 16 rats underwent viral infusions. Of the 16 virus rats, 8 rats were bilaterally injected with DREADDs virus AAV5-CaMKlla-hM4Di-mCherry (titer, 2.3×10^13^; Addgene http://addgene.org/50477) and 8 rats were bilaterally injected with AAV5-CaMKlla-mCherry control virus (titer, 2.3×10^13^; Addgene http://addgene.org/114469).

For the recording study (n = 9), rats were anesthetized with isoflurane (1-2% in O_2_) and mounted in a stereotaxic frame (Kopf, Tujunga CA). Prior to surgery Carprofen (5 mg/kg, SC) and sterile saline (3 mL, SC) were administered for analgesia and hydration, respectively. A pair of ground screws were implanted over cerebellum and a pair of anchor screws were implanted to ensure proper bonding to the 3D printed ring that was attached to the skull with C&B Metabond adhesive. Craniotomies were drilled over either HC and DLS contralaterally, HC and mPFC ipsilaterally, or DLS and mPFC ipsilaterally (Male coordinates HC: ± 3 ML -4 AP; DLS: ± 3.3 ML 0.7 AP; mPFC: ± 0.7 ML 2.8 AP; Female coordinates were adjusted by 93%). One 64-channel silicon probe was chronically implanted targeting each region, for a total of two probes in each rat. HC and DLS were always implanted with four shank 64-channel Cambridge Neurotech P-1 probes, while four mPFC rats were implanted with linear 64-channel Cambridge Neurotech H3 probes, one mPFC rat was implanted with two shank 64-channel Cambridge Neurotech H6 probe (R786 – simultaneous recordings from mPFC & HC), and one mPFC rat was implanted with four shank 64-channel Cambridge Neurotech P-1 probe (R787 – simultaneous recordings from mPFC & HC). Targeting for the left and right hemisphere was counterbalanced across rats accounting only for sex (Supplementary Fig. 1C, D, Supplementary Table 1).

The probes were mounted on custom-made 3D printed micro-drives for precise and independent adjustment of the vertical position. The probes were lowered to ∼1.5mm and ∼2.5 mm from skull surface for P-1 and H3/H6 probes respectively to ensure all recording sites were in the brain. The craniotomies were then filled with a sterile mixture of bone wax and mineral oil (3/1 by weight) and the micro-drives were adhered to the skull with Metabond. A 3D printed shell that was painted with conductive paint, head cap, and LED mount were attached to the skull ring, and the ground screw was attached to the inside of the shell via conductive epoxy.

For viral vector injections, rats went through the same pre-surgery protocol. Craniotomies were made bilaterally over mPFC (±0.7 ML 2.8 AP), and bilateral cannulae were lowered into depth -3.8 DV for females and -4 DV for males (Supplementary Fig. 7A). Rats were randomly assigned to receive either the active virus (CaMKlla-hM4Di(Gi)-mCherry; n = 8) or control virus (CaMKlla-mCherry; n = 8). 1μL of virus was delivered at a rate of 200 nL/min. The cannulae were slowly raised following a 15-minute wait period after infusion. The craniotomies were then filled with sterile bonewax. Finally, an LED attachment was adhered to the skull using C&B Metabond to facilitate video tracking during behavior.

At the completion of the surgery rats were given Baytril (10 mg/kg, SC), additional saline (20 ml/kg, SC), and Children’s Tylenol (0.8 mL, orally) for antibiotic, hydration and analgesia, respectively.

#### Left/Right/Alternation Task

The left/right/alternation task has been detailed in prior works.^1–5^ Briefly, in the left/right/alternation task (LRA), rats started from the Start of Maze region at the bottom of the track, and ran through a navigation sequence, in this version, that consisted of three low-cost choice points (Wall 1, Wall 2, Wall 3) to a high-cost choice point at the top of the track (Supplementary Fig. 1A). At the high-cost choice point the rats had to choose whether to proceed to the left or to the right, where each return track consisted of one feeder that automatically delivered one food pellet (45 mg, Test Diet, Richmond lN) if the choice was correct. The rats would then proceed back to the Start of Maze where one food pellet would be delivered if the current choice was correct. We refer to the region between Start of Maze and Choice Point as the central track. The walls of the maze were constructed using DUPLO LEGO bricks with vinyl flooring under the maze. Permanent walls had unique patterns but did not change throughout the experiment. The interior walls of the central path could change daily, but they remained constant within the day. For the recordings experiment, the maze was 108 cm × 163 cm with 25 cm wide tracks, and for the DREADDs virus study the maze was 94 cm × 100 cm with 13 cm wide tracks.

There were three different reward criteria that specified the correct choice on each lap, left reward (L) where food was delivered on the left return rail and Start of Maze, right reward (R) where food was delivered on the right return rail and Start of Maze, and alternation (A) where the rats had to alternate between the left and right return rails to receive reward at the chosen side and Start of Maze. For the left and right reward criteria a choice was considered correct if on each lap rats ran to the left and right sides of the track, respectively. For alternation, a choice was considered correct if on each lap rats alternated which side of the track they went (e.g., if prior choice was left the next choice had to be right and vice versa). If the starting rule was alternation, the rats always received a reward (both at side feeder and Start of Maze) on their first lap of the session.

At the beginning of the session, rats were placed on the track at the Start of Maze, and then were allowed to run continuously for 45 minutes. Rats were pre-trained to run in a loop. Each correct lap resulted in the automated delivery of two reward pellets, one at the chosen side and one at Start of Maze, thus with each correct lap, rats earned 90 mg of full-nutrition food reward (2 x 45 mg 5TUL TestDiet pellets). All pellets were unflavored and performance on the task constituted the sole opportunity for rats to earn food each day.

All behavioral events were operated through custom built Matlab software (Matlab v2015b, Mathworks) to track rat position, control task state, and trigger reward delivery. Video was collected via an overhead camera sampled at 30 Hz, tracking either the backpack LED (pre-op) or head-mounted LED (post-op).

Food delivery was via a Matlab-Arduino interface (Arduino UNO) sending TTL pulses to Med-Associate feeders (Fairfax VT).

#### Initial task training

Prior to training all rats were handled by the experimenter for 3 to 7 days until they were acclimated. The rats were then food restricted for 5 to 7 days and were offered 30 min of access to unflavored pellets (45 mg/pellet, TestDiet). After the rats were familiar and ate the available food, they were placed back on free feeding for 7 days to return them to their original free-feeding weight. Training started after food acclimation and weight rebound. The training took place before probe implantation for the recording study, and after the virus surgery for the manipulation study to ensure viral expression prior to the experimental phase. Behavioral tracking was done using backpacks with an LED for the recording rats, and a head-mounted LED for the virus rats. Rats were trained over three phases where each ranged from 5 to 15 days. During the first two phases there were no walls (low-cost choice points) within the central track, i.e., it was an open arena.

ln the first training phase, rats were trained to run in a loop starting from the Start of Maze. L and R rules were alternated each day. The incorrect (opposite to the session rule) arm was blocked such that rats were forced to turn congruent with the rule, and thus make the correct choice (e.g., if the reward contingency was L, the right arm was blocked). During this initial phase each feeder (side feeder and Start of Maze) delivered to 2 unflavored pellets/feeder (total of 4 pellets/lap, 180mg of food). Training continued until the rats were running continuously and were completing at least 50 laps/session for each reward contingency (left/right).

ln the second phase of training, blocks were removed, and alternation was introduced as a rule. During the rest of the training days rats experienced one pseudo-randomly chosen reward criterion per day.

Rats remained in this phase until they reached 80% correct for each reward criterion (left, right, or alternation) and ran ∼100 laps continuously (10-15 days).

ln the third phase of training, interior walls were introduced. All possible maze configurations within a given complexity (Fig. 2A) were randomly assigned to either be part of the training or the test set. Each day, a maze configuration from the training set was pseudo-randomly chosen without replacement, such that everyday had a unique maze configuration. LEGO DUPLO bricks were used to create the maze structure. Similar to training phase two, animals experienced a pseudo-randomly chosen single reward criterion. Rats remained in this phase until they reached 80% correct for each reward criterion (left, right, alternation) and ran ∼100 laps (5-7 days).

For the recording animals, surgery was conducted after training phase two. After the rats underwent probe implantation, they were reacclimated with the task starting from training phase two, during which the probes were lowered to their appropriate targets (7-12 days). After rats reached phase two criterion, they moved onto phase three, where along with the introduction of the maze walls, rats ran the task with the recording tether attached, familiarizing them with the process of performing the task under full task and recording conditions.

#### Experimental sequence

Once rats completed their training sequence and the probes were positioned at their appropriate targets, the rats immediately entered the experimental sequence, where all three reward rules were presented to the animal within a single session. Each 45 min session had two un-cued rule changes that occurred roughly 15 and 30 minutes into the session. A random amount of time between ± 2.2 mins was added or subtracted from the target switch times to reduce predictability. The rule sequence for a given session was pseudo-randomly drawn from one of six possibilities: LRA, LAR, RLA, RAL, ALR, ARL and maze configurations were pseudo-randomly chosen from the test set.

Recording animals (n = 9 rats) underwent a 16-day experimental sequence (1 session/day). Recordings were performed over these 16 days, totaling 144 recording sessions. Before the task start and after the task end rats sat on a flowerpot for 5 min to record baseline neural activity during quiet resting. The pre/post session recording data were processed but were otherwise not included in analysis.

The 16-day sequence was further broken up into two experimental sequences spanning 8 days. Each 8-day sequence consisted of 4 unique mazes that were randomly chosen without replacement along the complexity axis; however, each 8-day sequence included the simplest maze (AAA). Rats were run on the same maze configuration under the same task rules for 2 consecutive days (Supplementary Fig. 1B).

The same experimental protocol was used for the DREADDs rats; however, the experimental sequence was extended to 32-days (n_Sessions_ = 508), such that every maze configuration within the test set was used. Deschloroclozapine (DCZ; Hello Bio. Inc., Princeton NJ)^6^ was used to activate the DREADDs, or VEH (saline) was used as a control (Supplementary Fig. 7B). The active and control virus animals were paired such that each pair had the same exact same experimental sequence (mazes and switches). DCZ and VEH conditions were presented in a pseudorandomized order, controlling for first-order effects. Importantly, the 2^nd^ day-repeat of a given maze/rule configuration was always a saline washout day (Supplementary Fig. 7C). Every day the rats received subcutaneous (SC) injections of either DCZ, VEH, or saline 30 min prior to the start of the experiment. DCZ was dissolved in saline to a concentration of 0.05mg/mL and a single dosage of 0.1mg/kg of DCZ was used throughout the study. For VEH and saline a comparable volume was injected SC. All experimenters were blind to the identity of the solution (DCZ or VEH) injected on a given day.

#### Recordings

All recordings were collected at 30kHz sampling rate using a 256 channel lntan RHD recording system connected to two 64-channel lntan RHD headstages. An Arduino interface between Matlab and the lntan system facilitated time synchronization of neural data to ongoing behavior. In DLS and mPFC, throughout the 16-day recording sequence probes were periodically lowered (∼30μm intervals) to increase the likelihood of recording a diverse array of units. All movement of probes occurred after completion of day’s recordings leaving at least 23 hrs for probes to restabilize before the next recording session. After reaching the CA1 layer in dorsal HC during the training phase, the probes were not subsequently moved, unless there was visible spontaneous movement of the probe from one day to the next. In those cases, HC probes were either lowered or raised to ensure recordings from the CA1 cell layer. While we took appropriate efforts to record diverse ensemble sets of units across days, we are unable to claim that all recorded cells are unique. No attempts were made to identify recordings of the same cells across successive days.

Spikes were extracted and classified using Kilosort v1.0 for HC, and Kilosort v2.0 for DLS and mPFC using a custom pipeline. Kilosort v1.0 clustering for HC used previously published parameters tuned for HC (https://github.com/brendonw1/KilosortWrapper), and Kilosort v2.0 clustering for mPFC and DLS used default parameters.^7^ Automated sorting was followed by manual curation of the waveform clusters using Phy (https://github.com/cortex-lab/phy).^8^ Each silicon probe was analyzed independently and included the entire 45 min task data along with the 5 min pre/post recordings (∼55 min total recording time).

Prior to Kilosort processing, a median reference subtraction was performed across the 64 channels and a 600 Hz high-pass filter was used for spike identification. Collectively we identified 4292 single units from HC, 2293 single units from DLS, and 2609 single units from mPFC. All unit isolation and manual curation was performed blind to the session’s corresponding behavioral data.

HC units were separated into putative pyramidal cells and interneurons using their waveform characteristics and firing rates. DLS units were separated as phasic and nonphasic based on the distribution of inter-spike-intervals (ISIs).^9^ Phasic cells were those with at least 40% of all lSls greater than 2 seconds, which we take as putative medium spiny neurons (MSNs). Otherwise, the DLS cell was classified as a nonphasic firing cell. Following previous studies, if a cell that had a post-spike suppression index longer than 100 ms it was classified as tonic firing (putative tonically active cholinergic interneurons). If it did not meet either the MSN or TAN criterion, it was classified as high firing (putative high firing interneurons). Non-phasic (non-MSNs) cells were not analyzed further here.

#### DREADD validation

ln one additional rat (Male FBNF-1) we bilaterally transfused the dorsal mPFC (± 0.7 ML 2.8 AP -3 DV) with the same active inhibitory DREADDs used in the main study (AAV5-CaMKlla-hM4Di(Gi)-mCherry) (Supplementary Fig. 7B histology). Roughly six weeks after the viral injection, a single 64-channel Cambridge Neurotech H3 probe was chronically implanted into the infusion craniotomy on the right hemisphere and was lowered to 2.3 mm from skull surface. These surgeries used the same procedures outlined above. Over four days the probe was lowered to 4.13 mm (2.86 mm top of probe) from skull surface to span the volume around the DREADD infusion site (Supplementary Fig. 7B histology).

We performed two recording sessions, one with a DCZ^6^ injection and one with a VEH (saline) injection. The recordings were taken for ∼3hrs during quiet rest of the rat on a flowerpot. We first recorded a 30 min baseline period before drug injection. At 30 mins the rat was injected with either DCZ or VEH (1 session/drug), while the recordings were ongoing. The rat was then placed back on to the pot for another 2.5 hrs to establish a time course of drug effect. Multiunit spiking activity (18 isolated units taken collectively for VEH injection Supplementary Fig. 7B left panel, and 5 units for DCZ injection Supplementary Fig. 7B right panel) was binned into 30s bins to calculate the average firing rate over the 3-hour experiment.

#### Histology

Following the conclusion of all experiments, rats were overdosed with pentobarbital. Rats were perfused transcardially with phosphate-buffered saline and paraformaldehyde. For recording rats, probes were left in place for 3-5 hrs and were then carefully removed from the brain, leaving a visible track of their final position (Fig. 1B, Supplementary Fig. 1D). Brains were then extracted and post fixed in paraformaldehyde for 24 hrs before being transferred to a mixture of paraformaldehyde and sucrose for cryoprotection.

Coronal sections (40μm sections) surrounding the probe location were cut on a cryostat, mounted to slides. For recording rats, mounted sections were stained with cresyl violet for visualization. In two rats with DLS recordings we performed immunohistochemical staining for MOR1 on a subset of the tissue (anti-μ-Opioid Receptor gifted by Maureen Riedl). For virus rats, we used immunohistochemical procedures to amplify the endogenous -mCherry fluorophore. Mounted tissue was labeled with an anti-RFP antibody to visualize viral expression and with an anti-NeuN antibody to counter stain cell bodies. In both cases fluorescent secondary antibodies were used for visualization.

### Data Analysis

Most data analyses were performed using custom Matlab software (Matlab v2021a, Mathworks). Quantification of environment complexities were performed using the Networkx package for Python. Each subject was treated independently. No formal replication was performed, but our results agree with previously published work. For all analyses, data were pooled across all subjects (physiology or behavioral responses), but in all cases, we confirmed that reported findings were qualitatively consistent across rats and not driven by a single subject.

#### Quantification of maze complexity

Three different wall configurations were used to change the maze structure: *A:* two walls with open space in the middle, *B:* one wall with open space on the left, and *C*: one wall with open space on the right. Combinatorial combination with repetition of the three different wall types (A, B, C) created 27 unique mazes (3^3^ = 27) (Fig. 2A).

Graph theoretic measures have been previously used to quantify the complexity of an environment.^10–14^ To determine the structural complexity of a maze, we first discretized the inner navigation sequence into 15 bins. These bins corresponded to graph nodes, and each node was connected to another node if it could be reached by a cardinal action---i.e., an edge existed between the two nodes if there was no wall between the two bins. Starting from each node we created 15 generalized trees, with each root node being one of the 15 nodes of the discretized maze. The structural complexity of a tree graph is defined as the Shannon entropy of the distribution of the number of nodes at each level of the tree---e.g., the distribution of the number of nodes across all possible trees one step from the root node. The total structural complexity of the space was then defined as the total information content across all equivalent generalized trees that could be created from a given maze configuration.^10,11^ The distribution of the total information across mazes was then min-max normalized. This distribution had 3 distinct clusters denoted by the colored circles in Fig. 2A that were labeled as low (light green), mid (teal), and high complexity (dark blue) environments, respectively.

#### Behavior

Each lap (Start of Maze to Start of Maze) was identified as correct or error dependent on whether the rat chose the action congruent with the current contingency/rule and was subsequently rewarded. Percent correct for a given session was calculated based on the ratio between the number of correct laps and the total number of laps.

The percent correct for a given lap was calculated based on the ratio between the number of sessions in which that lap was correct and the total number of sessions, analyzed for each rat separately. Given that the sessions did not have equal number of laps and the switch times did not occur on the same lap, analyses were restricted to 10 laps pre- and 20 laps post-switch. Everything was computed for each switch independently and then averaged across the switches.

#### Behavioral change point

To identify laps where rats stabilized their behavior after rule switches, we used a modified version of the change point analysis developed by Gallistel et. al. (2004).^15^ Briefly, behavior in each lap was identified as a left or a right choice and was subsequently coded as either a 1 (left turn) or a 2 (right turn). Note that this differs slightly from prior implementations of this method which consider the correct/error nature of each choice as these methods can generate artificial change points when rats make an error on the first lap following a rule change (before they have any information about the change). This created a num. of Laps x 1 length vector. We then calculated the cumulative sum along this choice vector. Following Ref.^15^, we identified behavioral change points as those where the slope of the cumulative sum changed most abruptly. Computationally, this was done using the *ftndchangepts* function in Matlab with linear change point detection (*statistic = linear*) and minimum improvement in total residual set to 2 (*minthreshold = 2*) so as to not limit the number of change points that could be identified. After detection, change points in every session were visually confirmed by an experimenter blind to all information about maze configuration, task rules, and drug conditions.

#### Running speed

We computed the running speed as the change in *(x*, *y)* position *(dx*, *dy)* using an adaptive windowing of best-fit velocity vectors.^16^

#### Vicarious trial and error (VTE)

To quantify vicarious trial and error (VTE) behaviors at the choice point, we calculated the time integrated change in angular movement of the head (ldPhi), which captures both pause duration and the prominence of head re-orientation behaviors (Fig. 1F, G).^17,18^ ldPhi is calculated by the integral of the absolute value of the arctangent of the velocity over the duration that the rat is in the choice point.

Velocity *(dx*, *dy)*was calculated as the first derivative of the rat’s head position *(x*, *y)* using the Janabi-Sharifi and Marey (2010)^16^ method.

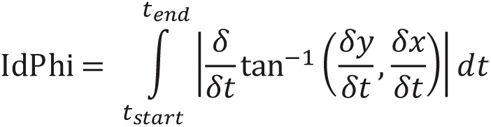

zldPhi was then computed as the z-scored ldPhi on a lap-by-lap basis for each session. In line with previous work, the distribution of the calculated zldPhi created a positively skewed bimodal distribution, where the prominent distribution (larger proportion of the overall data centered near 0) captured non-VTE events, and the smaller distribution tail captured VTE events. Based on this distribution, and similar to prior work, we identified a cutoff threshold between VTE or non-VTE laps as z = 0.25 (Supplementary Fig. 2D).^4,18^

#### Exploration

To calculate the amount of space explored, we binned the 2D space into ∼2cm bins. We then calculated the per session aggregate occupancy in the central path to create a 2D histogram (Fig. 3B, Supplementary Fig. 2l). The percentage of space explored was quantified as the total number of counts divided by the number of bins.

To classify a lap as explorative vs. non-explorative, we first found the median path through the central track. For each lap the trajectory through the central maze segment was resampled to 500 positional points to create a [num. of laps × 500] matrix for each *x* and *y* position. The median path was taken to be the median *(x*, *y)* location across laps for each interpolated sample. We then calculated the point-by-point distance between the resampled lap trajectory and the median path using the *dsearchn* function in Matlab. If more than 2.5% of the points in a given lap had distances that were above the 95% percentile of the overall distances across all laps, the lap in question was labeled as exploration.

#### Behavioral trajectory stereotypy

We calculated behavioral trajectory stereotypy from times that the rat was navigating through the central maze segment (central path) (Fig. 1H, l, Fig. 3F, G). Each central path trajectory was resampled to 500 positional points for each *x* and *y* position. For lap-by-lap stereotypy quantification, we found the reciprocal of the average Euclidean distance between lap *i* and lap *j* for each interpolated point.^19^ The stereotypy of a given lap *i* was calculated as the average of the trajectory similarity between lap *i* and ±5 laps around the diagonal of the stereotypy matrix (i.e., laps *i-2*, *i-1*, *i+1*, *i+2*-).

#### Maze linearization

The 2D maze was broken into 2 chunks: central path that spanned Start of Maze to Choice Point and return rails that spanned Choice Point to Start of Maze for each left and right turn (Supplementary Fig. 1A). To avoid any potential overlaps, the zone entry times for each lap were aligned such that the entry of one zone corresponded to the exit of another zone. We subsequently reflected the positional data that corresponded to the time when the rat was in the return rail zone to one side. Similar to above, we found both the median central path and the median return rail path and resampled them to 200 equidistant points. We then combined the resampled median central and return rail paths to form a single path across the whole track. This was done using the *polyshape* function in Matlab which created a fully enclosed polygon while respecting the overall structure of the input trajectory. This median track path was then resampled in 2D using the *interparc* function, which interpolates points along a 2D perimeter. All laps were then compared against this median template. The 2D *(x*, *y)* location of the actual position tracking was then reflected onto this median path using the *dsearchn* function, where the linearized 1D coordinate corresponded to the closest 2D position index, and thus binning the positional data in to 200 bins, which correspond to ∼2cm bins. All laps that were further than 20 cm away from the idealized path were excluded from further analysis. For each maze, we used the length of the median path through the entire maze (polygon perimeter) to calculate actual bin sizes in cm. For all analyses the binning was then down sampled from 200 to 150 bins resulting in 2.67 ± 0.03 cm bins for the simplest (AAA) maze and 2.89 ± 0.05 cm bins for the most complex (CBC) maze.

To account for any inconsistencies in linearized maze space due to different maze configurations, we identified the particular bins corresponding to relevant maze locations (e.g., Start of Maze, Wall, Choice Point) in each individual session. These identified bins were used for task bracketing analyses that depended on these designations.

#### Place field analysis

Tuning curves of HC pyramidal cells were calculated based on the linearized maze by first binning the space, and then generating the maps of bin spike counts over bin occupancy. Bins with occupancies less than 200 ms were omitted. A smoothed tuning curve (smoothing size: 2 bins) was constructed by convolving a Gaussian with the generated tuning curve. The smoothed tuning curves were then thresholded to 25% of their maximum firing rate, and place fields were defined as regions within the thresholded maps with an area greater than 15 cm and minimum firing rate of 1Hz. We then extracted the field centers and sizes using *regionprops* Matlab function on the binarized rate map.^20^

#### Hippocampal Theta cycle identification

The raw recordings taken from HC were decimated to 2 kHz to extract LFPs. Recordings were bandpass filtered with a Hilbert transform between 6-12 Hz to obtain the theta signal.^21–30^ The instantaneous phase and amplitude were estimated via the Hilbert transform. The channel to identify theta cycles from was chosen based on the probe channel that had maximal theta power. Given that Hilbert transform assumes the data is sinusoidal and because theta is asymmetric, using previous approaches^31^, we first found the peaks and troughs using the filtered signal. We then took these estimated peak and trough times in the raw signal and found the local minima (trough) and maxima (peak) around the estimated times. Theta cycles were limited to 80 to 200 ms. A single theta cycle was defined as peak-to-peak.

#### Bayesian decoding and sequence analysis

We used Bayesian decoding to calculate ensemble neural representations at both behavioral time-scales and theta-timescales.^32^ Briefly, this method computes the posterior probability of the ensemble representing each position in space given spike counts from each single unit in the ensemble. Posterior distributions were normalized to sum to one. Probability of an animal’s position *x* given all observed spikes from recorded units can be operationalized by:

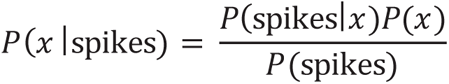

where *x* is the set of all linear positions on the track, binned in to ∼3cm bins. This method assumes that every cell is independent from each other and follows a Poisson process.^32^ For decoding of behavioral sequences, as in Fig. 4C, we decoded spiking in 120 ms time windows using a uniform spatial prior.

Bayesian decoding was 5-fold cross-validated. To implement k-fold cross-validation we divided the data into *k* equally sized subsamples. Of the k-subsamples, *k-* 1 subsamples were used as the training data and the remaining single subsample was used as the test data. The spikes and position were masked for the duration of the test data to create a *P*(spikes|*x*) likelihood function from the training data (created using the *k -* 1 subsamples). The posterior probability was calculated using the above equation for the test data (i.e., *P*(spikes) and *P*(*x*) were distributions based on the single subsample).

For all theta-timescale decoding analyses we used two criteria: 1) minimum of 5 cells had to be spiking in the given theta cycle, and 2) the running speed of the rat had to be faster than 2 cm/s. To examine the spatial decoding in theta cycles (Fig. 4l), we decoded spiking in each half of the theta cycle — peak to trough and trough to subsequent peak — to obtain a probability distribution across the linearized maze. The obtained posterior probability distribution was subsequently aligned to the rat’s position (± 50 cm around the rat). Posterior probabilities obtained from the two halves of theta were subtracted from each other (2^nd^ half – 1^st^ half) to obtain the local vs non-local decoding.^33,34^ The null distribution was obtained by shuffling (n = 1000) the identity of the decoding, 1^st^ vs 2^nd^ half (red line in Fig. 4l).

To identify sequential structure of theta cycles, we adapted two widely used measures: 1) theta sequence score,^35,36^ and 2) theta sequence slope.^36–38^ ln both cases we used Bayesian decoding methods with time bins defined by 1/6 of a theta cycle measure (measure 1) or 20ms bins (measure 2). In the first method, we used weighted correlation coefficients between time and the decoding posterior *r*(*x*, *t*|*P*). The posterior probability matrix *P* [num. of spatial bins × num. of time points] was used as weights. In a general sense, correlation coefficient is simply the ratio of the covariances of the two variables, normalized by the square root of their variances. Thus, in this case, the weighted means for location *x* and time *t* can be written as:

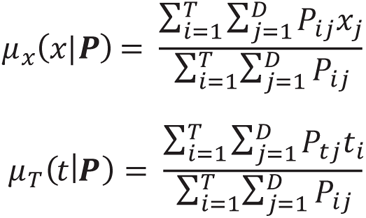

where *P_ij_* is the probability at time *i* and spatial bin *j*. The weighted covariance therefore can be written as:

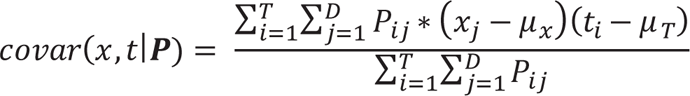

Finally, the weighted correlation can be calculated by:

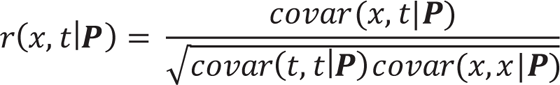

Weighted correlation for each reconstructed theta sequence was calculated from the decoded probabilities of positions 50 cm ahead and 20 cm behind the animal’s current position, and decoding was performed for time windows corresponding to *^rr^_/_*_6_ phase intervals of each theta cycle. Intuitively, a sequential structure sweeping along the animal’s running direction would yield a large positive correlation, whereas lack of structure would be close to 0.

ln the second method we computed theta sequence slope by fitting a line that yielded maximum decoded probability within a ± 20 cm window around the rat. For a given line with slope *v* and intercept *p*, the average likelihood *R*(*v*, *p*), that the decoded position is located within a distance *d* of that line can be written as:

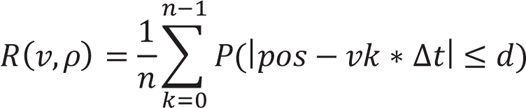

where *k* is the temporal bin of the posterior probability matrix and *Mt* is the moving step of the decoding window, in this case 20 ms. The value of *d* was set to be 10 cm around the rat. In line with prior research, we densely sampled the parameter space for *v* to find the value that maximized *R*.

To calculate the significance of either theta sequences (both sequence score and sequence slope), we circularly shifted the space-bins of the posterior probability distribution (n = 1000) for each time point. As previously described this shuffle conserves the structure of decoded probabilities within a bin but randomizes association between bins. A theta sequence was significant if its score was above the 95^th^ or below the 5^th^ percentile of the shuffled distribution to account for both forward and reverse sequences.^35,36,38^

#### Correlation between theta metrics and behavior

Three behavioral metrics were correlated to theta sequence score and theta sequence slope (Fig. 5, Supplementary Fig. 4). For exploration and stereotypy correlations, only theta sequences within the central path were considered. For zldPhi correlations, only theta sequences at the choice point were considered. For each session we performed a lap-by-lap correlation between the median theta sequence metric and the behavioral metric (Fig. 5A, C, E & Supplementary Fig. 4A, C, E). For binarized exploration vs. no exploration or VTE vs. no VTE, for each session we calculated the median theta score of explorative laps, VTE laps, non-explorative laps, and non-VTE laps, thus the distributions reflect session means (Fig. 5B, D, Supplementary Fig. 4B, D).

#### Linearized firing rate by lap

To construct the firing rate maps in Fig. 6A, we created rate maps for every lap and every DLS cell as a function of the linear location on the maze (binned into ∼3cm bins) resulting in a [laps × num. of spatial bins × num. of cells] matrix (35.8 ± 2.8 SEM number of single units/session). For each session we took the 25 laps from the start of the session and 25 laps following a rule switch (overall rats ran 45.7 ± 12.9 number of laps/rule block), and averaged firing rates across sessions for each maze complexity (low, mid, and high).

#### Task-bracketing index

Following previous research, we calculated the task bracketing index by subtracting the mean normalized firing rate in the central path from the mean normalized firing rate at the Start of Maze for each cell (Fig. 6B).^19,39–42^ Zones were defined based on entry and exit times for each session. Positive task bracketing indices indicate relatively strong firing in the Start of Maze compared to the subsequent navigation through the central path, while indices near 0 indicate that there is little to no difference in firing rate at the start compared to the rest of the maze. Given the observed firing rates of DLS cells across the linearized maze, we replicated the analysis using the start and end positions as Start of Maze and Choice point, and this analysis also yielded qualitatively similar results.

#### Spatial cross correlation of ensemble activity

To investigate how spatial representations were similar to each other across the linearized maze we calculated the spatial cross correlation of ensemble activity (Fig. 6D, Supplementary Fig. 5C-F). We correlated neural rate maps between pairs of spatial bins. For each cell, we calculated the firing rate in each of the spatial bins throughout the entire maze. For each pair of spatial bins, we computed the correlation between the normalized firing rate vectors for those bins. Therefore, if representations are local (e.g., as is in HC Supplementary Fig. 5C) the spatial cross correlation would appear as highly restricted around the diagonal of the matrix.

#### Stepwise regression between DLS firing and behavior

To account for linear relationships between behavioral variables (velocity and head direction), we used a stepwise linear regression procedure to relate firing of DLS cells to independent behavioral variables. We performed a sequential and single variable stepwise linear regression to progressively remove the influence of a given behavioral variable on neural firing. Starting from fitting both variables to DLS firing rates, after each step regression subsequent behaviors were fit to the residual firing rates.

For each lap, velocity in a given linearized spatial bin was averaged to create a lap by bin matrix. Given that we did not have access to exact head reorientation behaviors, we used a proxy, ldPhi, which was independently calculated for each linearized spatial bin. For single variable regressions, each behavior of interest was regressed out separately. For the stepwise linear regression, velocity was regressed out first followed by ldPhi.

#### Population correlation of ensemble activity

Following previous research, to measure the similarity between population activity across laps, we correlated neural rate maps between pairs of laps.^1,4^ For each cell, we calculated the firing rate in each of the 5 zone bins (Supplementary Fig. 1A) along the central stem (Start of Maze, Wall 1, Wall 2, Wall 3, Choice point). For each pair of laps, we computed the correlation between the normalized firing rate vectors for the lap. We averaged windows of 10 laps prior and 20 laps post rule switch across sessions to generate switch-aligned average correlation matrices for mPFC, HC, and DLS (Fig. 7A).

#### Ensemble population activity change point

To calculate the population activity change point we used the individual session population correlations. To determine where there are abrupt changes to the population correlation matrix, we used hierarchical clustering to identify reliable neural representations. Briefly, this method seeks to cluster data in an input matrix by constructing a hierarchical cluster tree. Unlike k-means clustering, because the linking tree is fixed, the starting point is not random.

We first created the linking tree for the population correlation matrix using the *linkage* function in Matlab and set the method to minimize the inner Euclidean distance between elements (*method = ward*). Subsequently, we performed agglomerative clustering of the created hierarchical cluster tree (linkage tree) by using *cluster* function in Matlab. Because the clustering depends on the number of allowed clusters, we followed previous work and repeated our analysis multiple times allowing the maximum clusters to range from 3 to 7.^1,4^ We then applied a change point analysis similar to that for detecting behavioral changes,^15^ but based on cluster identity number instead of choice identity. For each cluster size and session, we cumulatively summed the output cluster lDs, and used the *ftndchangepts* function in Matlab to find the laps in which there was a significant change in the cumulative sum of cluster IDs based on slope and mean (*statistic = linear*). Intuitively, this change point reflects the moment in which neural ensemble data shifted from one consistent representation to another. For each cluster size and session, to calculate the null distribution for lap change points, we shuffled (n_shuffles_ = 75) the cluster lDs across laps and found the change point of the shuffled data.

We calculated the distribution of change points aligned to change points as the difference between all the change points identified by the clustering in a given switch block to the block’s switch lap. The minimum distance was set as the minimum of the above distribution.

#### mPFC activity prediction of theta score

We used 5-fold cross-validated Bayesian decoding to decode whether ensemble activity in mPFC predicted hippocampal theta sequence score. For each theta cycle, we calculated the spiking activity of mPFC ensembles. We only took theta cycles in which there were at least 5 mPFC and HC cells active ensuring that we were only using theta cycles in which we had a valid theta sequence score.

Bayesian decoding requires the expected firing of a cell given the variable of interest to create the likelihood function. We defined our variable of interest as the theta sequence score (15 bins), such that our likelihood function was the expected firing of mPFC cells given binned theta sequence score. Similar to before, we assumed a uniform prior over theta sequence scores. Subsequently, for each theta cycle, we identified the discretized bin of its respective theta sequence score and found the decoded posterior probability for that particular bin — the probability of identifying the correct theta sequence score bin for the particular theta cycles, given a set of mPFC activity. Similar to Bayesian decoding of position, to cross-validate our data, we equally partitioned our data into 5 subsamples. 4 subsamples were used as the training data to create the above likelihood function, and the other remaining subsample was used to calculate the posterior.

We used three different shuffle methods to create null distributions:^22^ 1) shuffled mPFC spike timings, which breaks any relationships within mPFC and/or between mPFC and HC, 2) across session shuffled theta sequence scores, which preserves the mPFC representations, but breaks any moment-by-moment relationship between mPFC and HC, 3) within-lap shuffled theta sequence scores, which breaks the relationship between mPFC and HC within a given lap, but preserves longer behaviorally-relevant timescale relationships.

ln the first method, we shuffled mPFC spike times by circularly shifting the full spike train of each cell by a random amount ranging from 1 min to 45 mins to disassociate neural spiking from variable of interest. In the second method, we shuffled all theta sequence scores within a session by randomly permuting the vector of theta sequence scores to disassociate theta sequence scores from neural spiking in mPFC. Finally, in the third method, we isolated theta sequence scores for each lap, and similarly, randomly permuted theta sequence scores within a given lap. For each shuffle we used 5-fold Bayesian decoding to create null distributions. All shuffles were performed 5 times and the mean value was taken across the shuffles as the respective null value.

Per lap mPFC posterior probability around rule switches were computed by first restricting the data (theta cycles and mPFC activity) to start of maze to side feeder entry and then decoding sequence scores for those theta cycles given restricted mPFC activity. Similar to what was done above, we shuffled the restricted mPFC spike times by circularly shifting them and using the shuffled activity to predict the restricted HC sequence score.

#### Correlation between mPFC prediction of theta score and behavior

Similar to the analysis done relating theta sequence scores to exploration and zldPhi for a given pass through the central path for exploration or a pass through the choice point for zldPhi, we calculated the median posterior probability and correlated that to each metric independently. For each lap, we calculated the correlation between mean deviation through the central path and zldPhi at the choice point and the median posterior probability of mPFC prediction of HC theta sequence score. We also did the same calculation for null distributions created by the shuffles.

#### Lap peri-event time histograms of mPFC activity

Full lap switch-aligned mPFC activity were computed using standard methods. For each lap (Start of Maze to Start of Maze), we totaled the spike count and divided by the total lap time to compute average firing rate in each lap. We then normalized mPFC activity across laps for each cell. To compute mPFC activity in different zones, we restricted per lap mPFC activity to either side feeder times (from Feeder entry to Feeder exit) or to the central path (Start of Maze exit to Choice Point entry) and computed spike counts for each of the zones and divided it by the time spent in each zone to calculate firing rates.

Similarly, we normalized mPFC activity across laps for each cell.

To examine mPFC activity for correct and error trials, we labeled each lap as either correct or error based on choice and used the normalized full lap mPFC activity for each correct and error trial across all sessions to create the full distributions.

### Quantification and statistical analysis

#### Statistical analysis

Data analyses were performed using custom MATLAB (MATLAB 2021a, Mathworks) functions. For comparison across environmental complexity we used mixed effects ANOVAs setting rat as a nested variable for either %Correct or proportion of VTE events and rat as a random factor when evaluating behavioral data and a Kruskal-Wallis test when evaluating physiology. For comparisons between pairs of groups, we used a Wilcoxon rank-sum test. When relating exploratory and procedural behavioral parameters to HC theta sequence score, we used one sample t-tests. For comparison of behavioral responses to different drug conditions, we used mixed effects ANOVAs and set rat as a nested variable and as a random factor. Post-hoc analyses between conditions was done using rank-sum with Bonferroni corrections. For comparison of behavioral responses in DREADD experiments across days, we used a Wilcoxon signed-rank test and mixed effects ANOVAs with rat as a nested variable in the behavior of interest and as a random factor. All assessments of behavior and physiologic responses across laps around a rule switch was done with a one-way repeated measures ANOVA with pairwise post-hoc via rank-sum with Bonferroni corrections. All significance was assessed at a = 0.05.

---

**Supplementary Figure 1:**
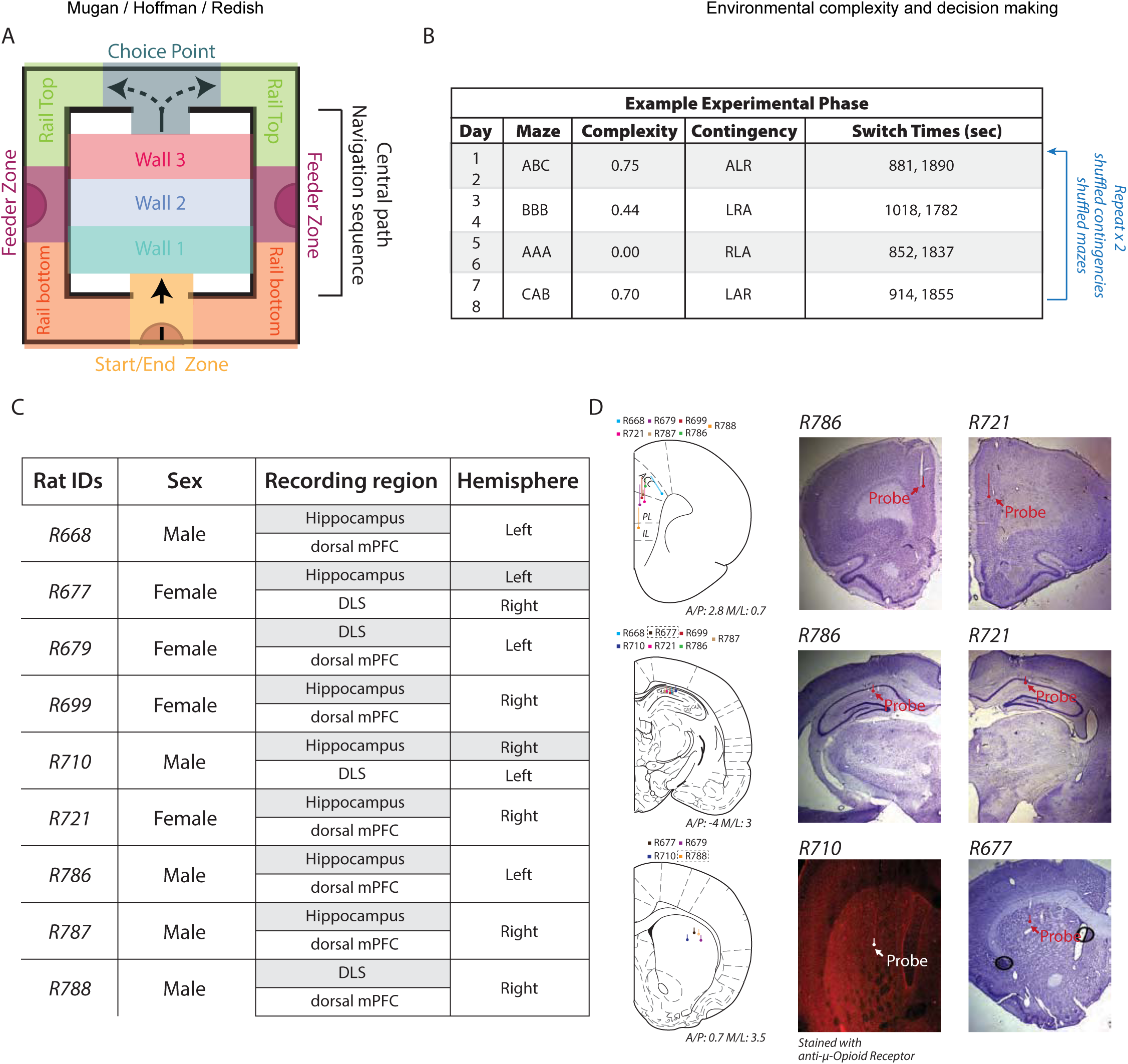
Task design and recordings, Related to Figure 1. (A) The left/right/alternation maze. The colored blocks indicate the 11 zones the maze was broken into. (B) An example 8-day sequence of the experimental paradigm. A set of 4 mazes were selected across the complexity space. Rats were run on the same maze (i.e., same central path configuration) with the same rules for 2 subsequent days. This 8-day sequence was repeated 2 times with randomized mazes and rule sequences. (C) Breakdown of the sex and probe targeting across rats. (D) Left: Silicon probe recording locations in prefrontal cortex (top row), hippocampus (middle row) and dorsolateral striatum (bottom row). Circles indicate the termination of the recording sites, and the extending line indicates the length of the recording sites. For 2 rats (outlined), no histology was available and so identified locations are estimates based on targeting coordinates, turning notes, and physiologic landmarks. Right: Histological examples of recording sites from prefrontal cortex, hippocampus, and dorsolateral striatum.

**Supplementary Figure 2:**
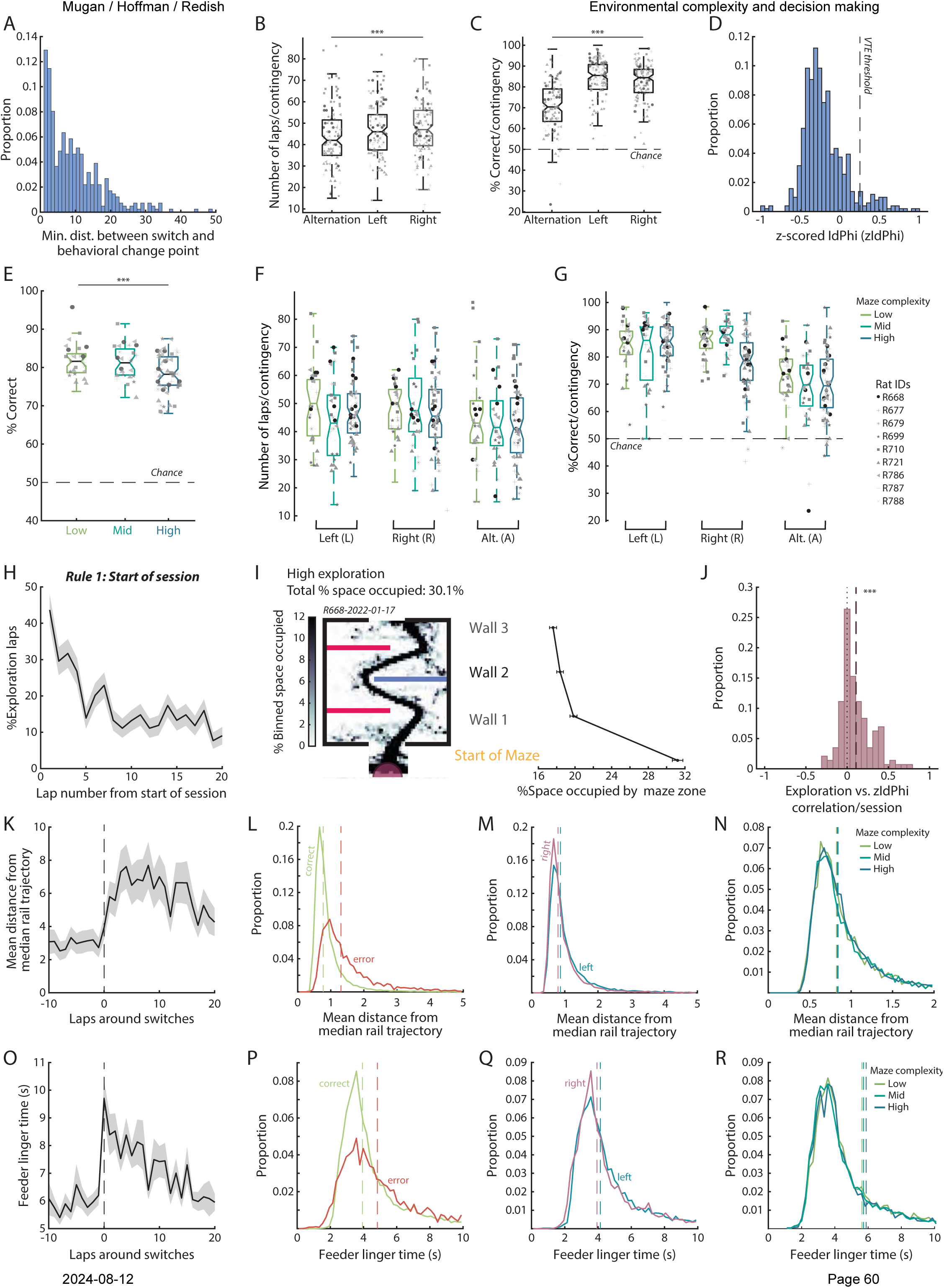
Behavior on the left/right/alternation task, Related to Figure 1 and Figure 2. **(A)** Distribution of minimum distance between detected behavioral change points to the corresponding task rule switch across all sessions. **(B)** Number of laps run per contingency/rule identity (Mixed effects ANOVA F(18) = 3.61, p < 0.001). **(C)** Rat performance for each contingency/rule (Mixed effects ANOVA F(18) = 13.47, p < 0.001). **(D)** Distribution of all zldPhi values. zldPhi was a positively skewed bimodal distribution, and in line with prior research, we set the threshold of VTE to be around the cutoff between the two peaks (0.25). **(E)** Rat performance in low, mid, and high complexity environments (Mixed effects ANOVA F(18) = 2.35, p = 0.0033). **(F)** Number of laps run per contingency/rule across different complexity groups (Mixed effects ANOVA laps/rule*complexity interaction F(36) = 2.85, p < 0.001). **(G)** Rat performance for each contingency/rule across the different complexity groups (Mixed effects ANOVA performance/rule*complexity interaction F(36) = 2.16, p < 0.001). **(H)** Proportion of exploratory laps aligned to the start of the session (initial rule block). The black line indicates mean, and the shaded gray region ± SEM. **(I)** *Left*: An example of high exploration of the start of maze and central maze segment (central path). *Right*: Quantification across all sessions of the percent space occupied in each maze region (see Supplementary Fig. 1A for maze zone designations). The dots indicate mean and the error bars ± SEM. **(J)** Distribution of correlations between zldPhi and exploration amount (t-test t(143) = 6.86, p < 0.001). Exploration amount was calculated as the lap’s average deviation from the median path through the central maze segment. The vertical dashed line indicates the mean. **(K)** Mean point-by-point difference between path taken in the return rail (choice point exit to start of maze entry, see Supplementary Fig. 1A for designations) and the overall median lap (mean deviation from median path) aligned to rule switches. Representation as in (H). Intuitively, greater average deviance indicates less stereotypy. **(L)** Distribution of side rail mean path deviation for correct (green) and error (red) trials. Dashed lines indicate the medians. **(M)** Distribution of side rail mean path deviation for left (blue) and right (purple) trials. Dashed lines indicate the medians. **(N)** Distribution of side rail mean path deviation for the different maze complexity groups. Dashed lines indicate medians. **(O)** Time spent in either the left or right feeder zones aligned to rule switches. Representation as in (H). **(P)** Distribution of time spent in either the left or the right feeder for correct (green) or error (red) trials. Representation as in (L). **(Q)** Distribution of time spent in the left (blue) or the right (purple) feeder. Representation as in (M). **(R)** Distribution of time spent in either the left or right feeder for the different maze complexity groups. Representation as in (N).

**Supplementary Figure 3:**
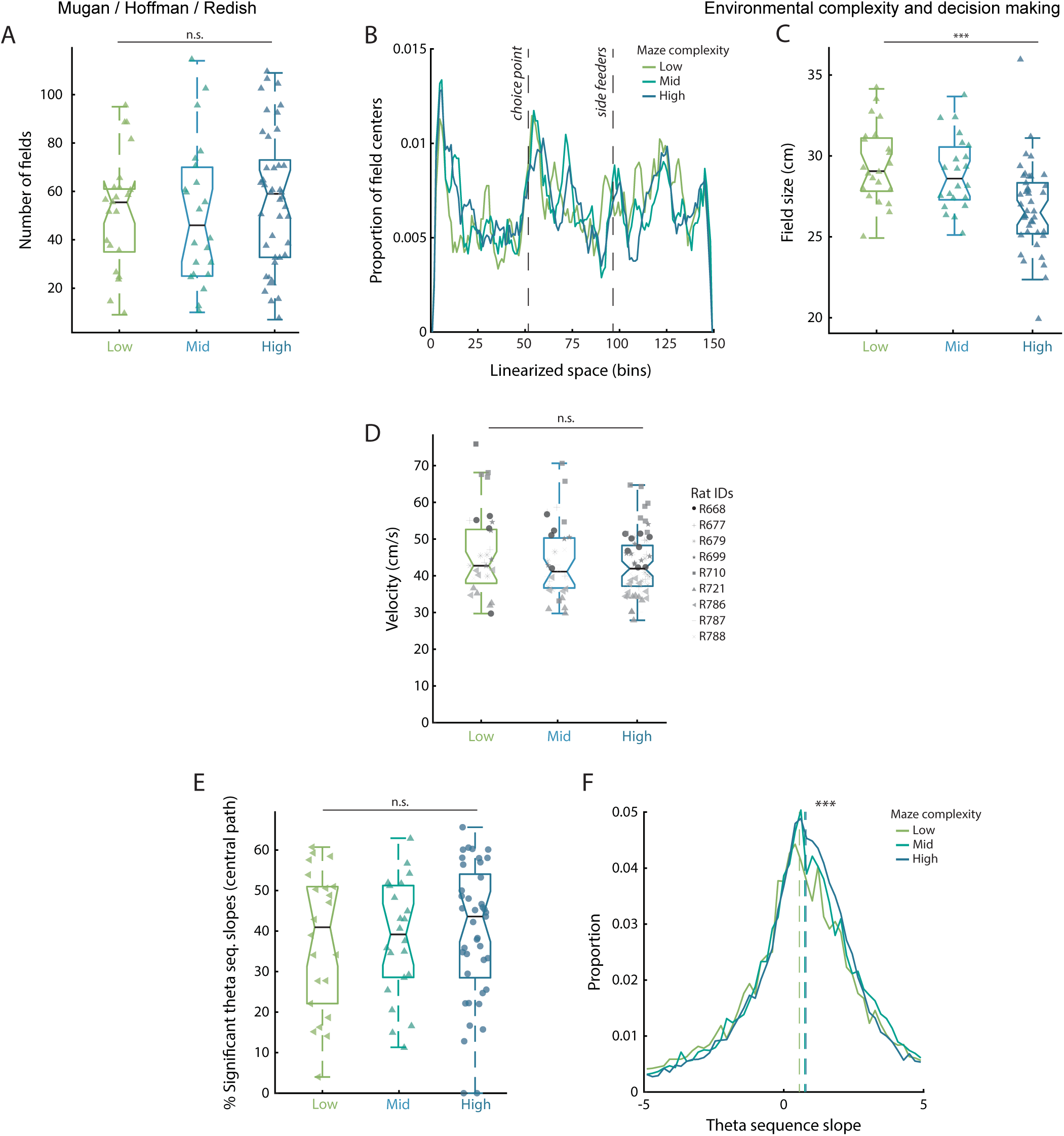
Basic hippocampal physiology and theta sequence slopes in dilferently-complex environments, Related to Figure 4. **(A)** Total number of HC place fields broken down by complexity. There was no significant difference in the number of fields detected (One-way ANOVA F(2) = 0.53, p = 0.59). **(B)** Distribution of HC place field centers across the linearized maze (2.67 ± 0.03 cm bins for the simplest maze and 2.89 ± 0.05 cm bins for the most complex maze)). The distribution of HC place field centers is largely uniform with a slight overrepresentation at the start and choice point. However, the complexity and structure of the maze did not affect the distribution of the place fields. **(C)** HC place field sizes broken down by complexity. In more complex environments fields were smaller (One-way ANOVA F(2) = 9.91, p = 0.0001). **(D)** Distribution of speeds through the central path for different maze complexities (Mixed effects ANOVA F(18) = 1.29, p = 0.20). **(E, F)** Same as Fig. 4F, G. but using theta sequence slopes instead of theta sequence scores. (E) Average percent of significant theta sequence slopes per session. Similar to Fig. 3F we did not find a significant difference in the proportion of theta sequence slopes that were identified as significant (see Methods) (Mixed effects ANOVA F(12) = 1.12, p = 0.36). (F) Distribution of theta sequence slopes of the different maze complexity groups. Similar to Fig. 4G we found that complex environments favored steeper theta sequences (Kruskal-Wallis H(2) = 66.67, p < 0.001).

**Supplementary Figure 4:**
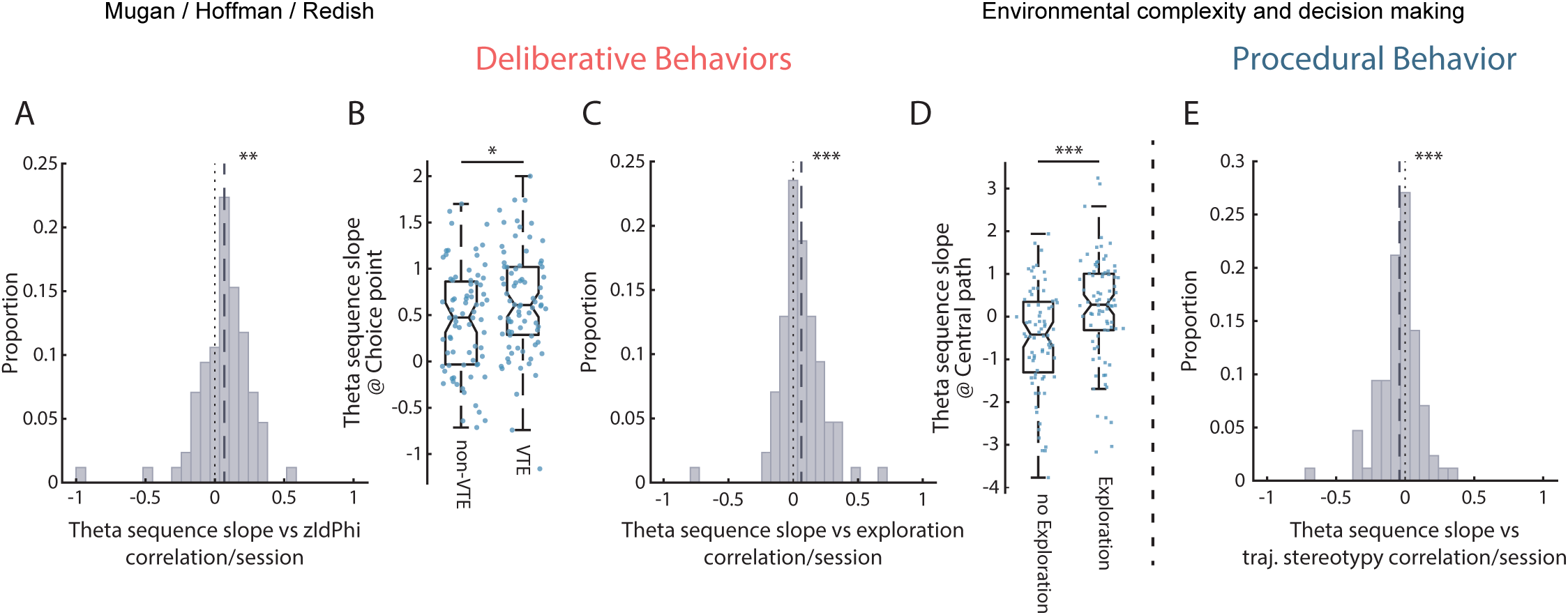
Exploration as a deliberative behavior and the relation between theta sequence slopes and deliberative behaviors, Related to Figure 5. **(A-E)** Same analysis as Fig. 5A-E but for theta sequence slopes. (A) Distribution of correlation between theta sequence slopes and zldPhi (t-test(80) = -3.07, p = 0.0015). (B) Theta sequence slopes for VTE and non-VTE laps (Kruskal-Wallis Test H(1) = 3.82, p = 0.05). (C) Distribution of correlation between theta sequence slopes and exploration amount (t-test(84) = -3.29, p = 0.0015). (D) Theta sequence slopes for exploratory and non-exploratory laps (Kruskal-Wallis Test H(1) = 12, p = 0.0005). (E) Distribution of correlation between theta sequence slope and path stereotypy through the central maze segment (t-test (84) = 2.56, p = 0.0122). For panels A, C, E the vertical line indicates the means.

**Supplementary Figure 5:**
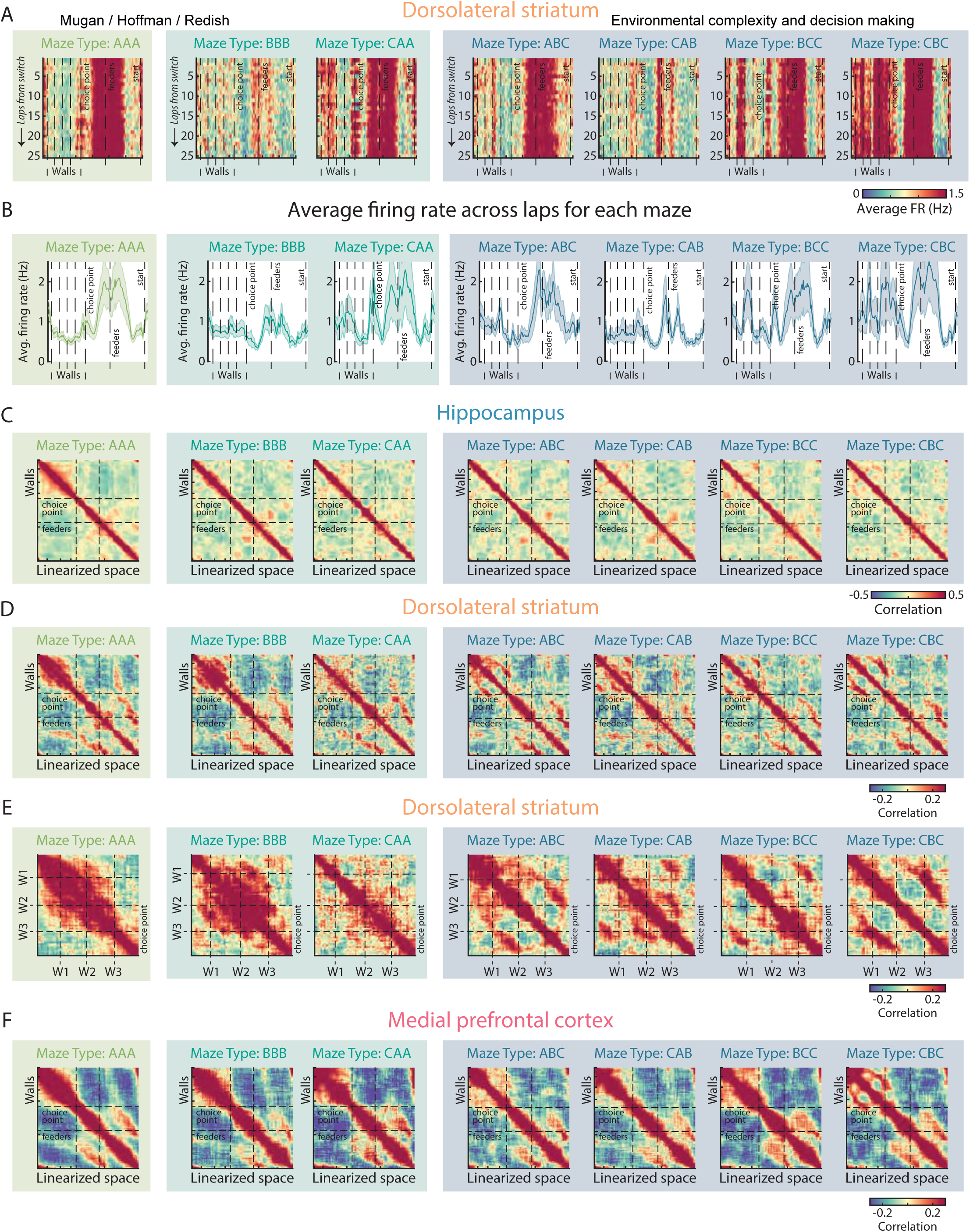
Spatial cross correlation of HC and DLS, Related to Figure 6. **(A)** Display as in Fig. 6A top row. Average firing rate of putative MSNs over the linearized track for each tested maze configuration. Black dashed lines indicate the wall locations (A, B, or C) for a given maze configuration. Bin size as in Fig. 6A. (B) Display as in Fig. 6A bottom row. Average firing rates across laps over the linearized track for each tested maze configuration. Solid lines indicate the mean, and the shading ± SEM. Black dashed lines indicate the internal wall locations projected onto the linearized maze. (C) Display as in Fig. 6D. Spatial cross correlation of HC ensemble activity for each tested maze configuration. The dashed lines indicate the choice point and feeder sites. HC shows largely local representations throughout the linearized maze (Bin size as in Fig. 5A). **(D)** Display as in Fig. 6D. Spatial cross correlation of DLS ensemble activity for each tested maze zone. **(E)** Display as in (D). Spatial cross correlation of DLS ensemble activity in the central segment for each tested maze. Black dashed lines indicate the internal wall locations projected onto the linearized maze (Bin sizes as in Fig. 6D). **(F)** Display as in (C, D). Spatial cross correlation of mPFC ensemble activity for each tested maze configuration.

**Supplementary Figure 6:**
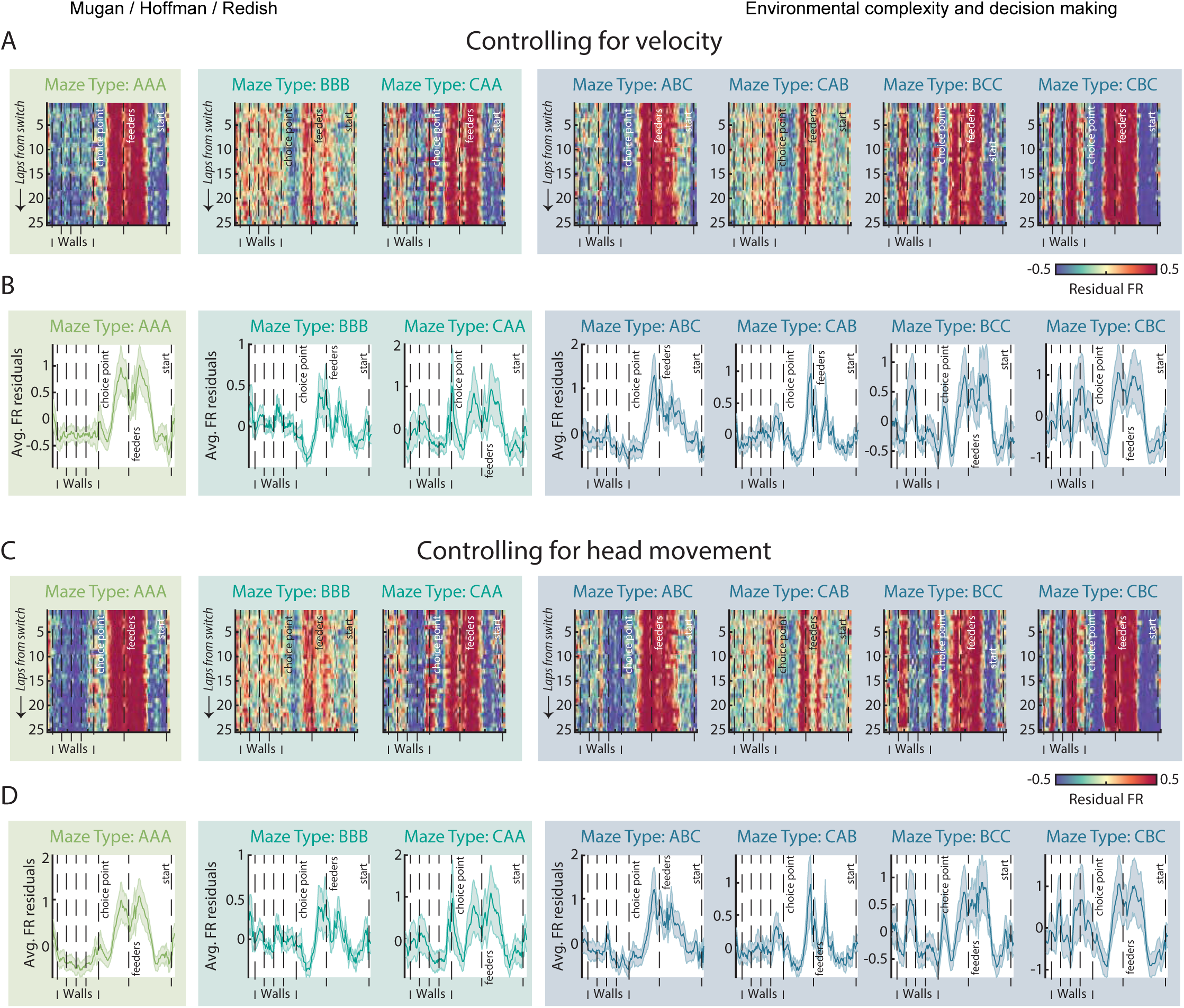
DLS firing rate residuals after controlling for velocity and head movement, Related to Figure 6. **(A)** Display as in Fig. 5A top row. Average residual firing rate of DLS neurons after controlling for velocity across the linearized maze for each tested maze. Black dashed lines indicate the wall locations, choice point, and feeder locations. Bin size as in Fig. 6A. **(B)** Display as in Fig. 6A bottom row. Average residual firing rate of DLS neurons after controlling for velocity across laps for each tested maze configuration. Solid line indicates mean, and the shading ± SEM. Black dashed lines indicate the wall locations, choice point, and feeder locations. **(C, D)** Residual firing rate of DLS neurons across the linearized maze for each tested maze configuration when controlling for ldPhi values in a given linearized location. (C) Representation as in (A) and (D) Representation as in (B).

**Supplementary Figure 7:**
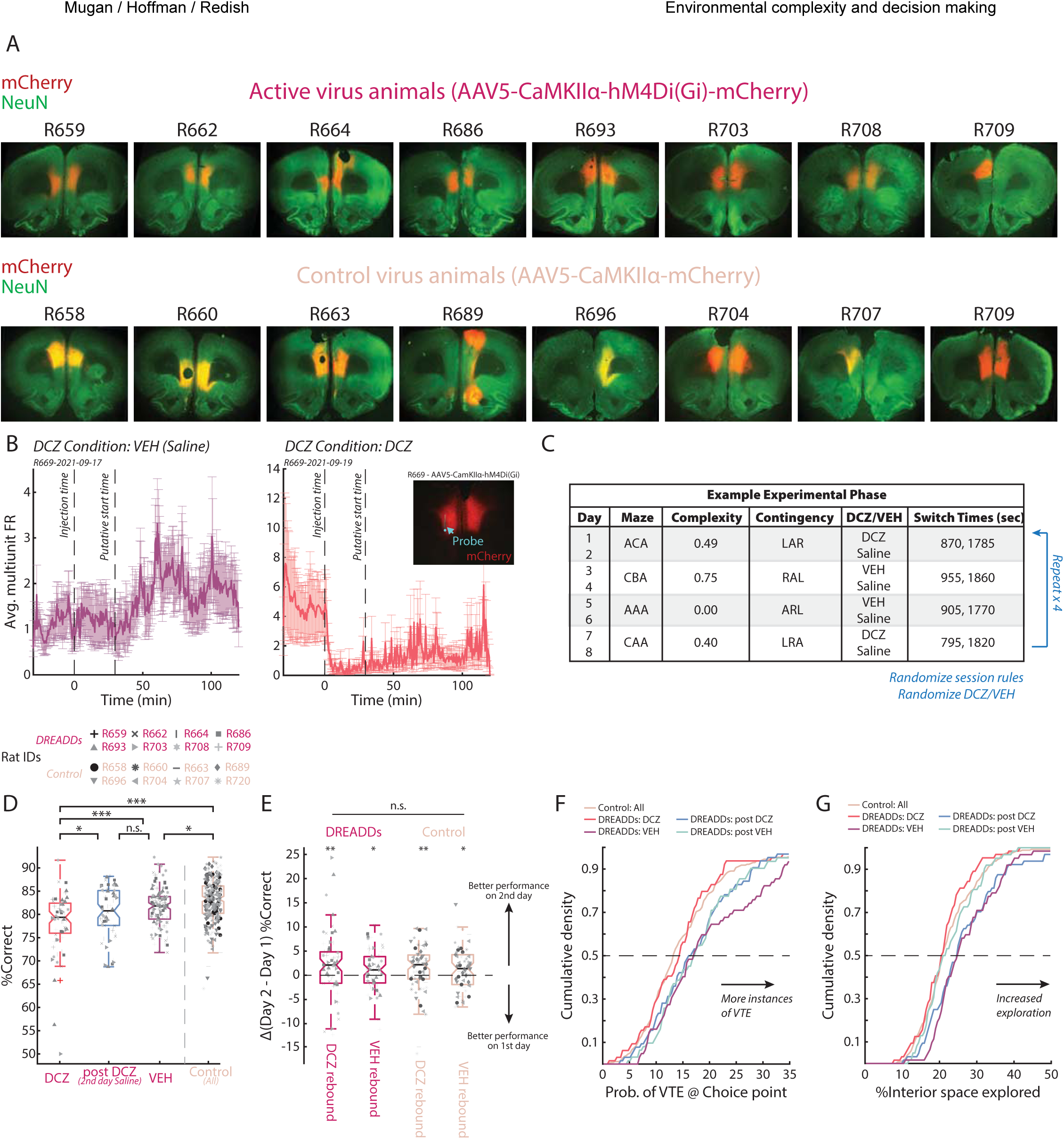
Histology and behavioral elfects of mPFC inactivation with DREADDs, Related to Figure 8. **(A)** All histological sections from virus transfected rats included in the study. Top row shows the histological sections from rats transfected with the active DREADDs virus (AAV5-CaMKlla-hM4Di(Gi)-mCherry), and the bottom row shows the histological sections from rats transfected with the control virus (AAV5-CAMKlla-mCherry). Note that in most cases expression is bilateral and localized to dorsal mPFC (anterior cingulate and prelimbic cortices). **(B)** Proof of concept for DREADDs and DCZ inactivation (n_Rats_ = 1, n_Sessions_ = 2). Average multiunit firing over 3hrs while the rat was on a flowerpot to measure the changes to baseline neural activity under the VEH (Left) and DCZ (Right) conditions. The vertical lines indicate the time of injection and the putative experiment start time (∼30 mins after injection). Inset shows the histology for the DREADDs expression and probe placement. **(C)** An example 8-day sequence of the experimental paradigm for prefrontal manipulations. A total of 14 different mazes were used and selected from different variations of the maze inserts to span the complexity across the entire experimental phase. 4 unique mazes that spanned the complexity space were used across the 8-day sequence. Rats were run on the same maze (i.e., same central path configuration) with the same contingencies for 2 consecutive days, in which the second day was always a saline washout day. This 8- day sequence was repeated 4 times with randomized mazes, contingencies, and drug conditions. **(D)** Distribution of performance on the task (% correct) for DREADDs virus rats under DCZ, post DCZ Saline, and VEH, as well as all control virus animals (One-way ANOVA F(3) = 18.31, p < 0.001; HSD Post-hoc DCZ vs all others p < 0.05, post DCZ Saline vs VEH p = 0.43, control vs all others p < 0.05). **(E)** Difference in performance between 1^st^ day manipulation and 2^nd^ day rebound (Day 2 – Day 1) for DREADDs and control virus animals with respect to drug condition. There was no significant effect of virus, and all the rats significantly increased their performance on the repeat day (2^nd^ day) (ANOVA virus F(1) = 0.35, p = 0.55; Wilcoxon signed rank DREADDs: rebound DCZ z = 2.38, p = 0.0088; DREADDs: rebound VEH z = 1.65, p = 0.049; Control: rebound DCZ z = 2.37, p = 0.0088; Control: rebound VEH z = 1.91, p = 0.028). **(F, G)** Same as Fig. 8D, F. (F) Distribution of proportion of VTE events/session for different drug conditions (G) Distribution of the occupancy of the central segment of the maze/session for different drug conditions.

**Supplementary Fig. 8:**
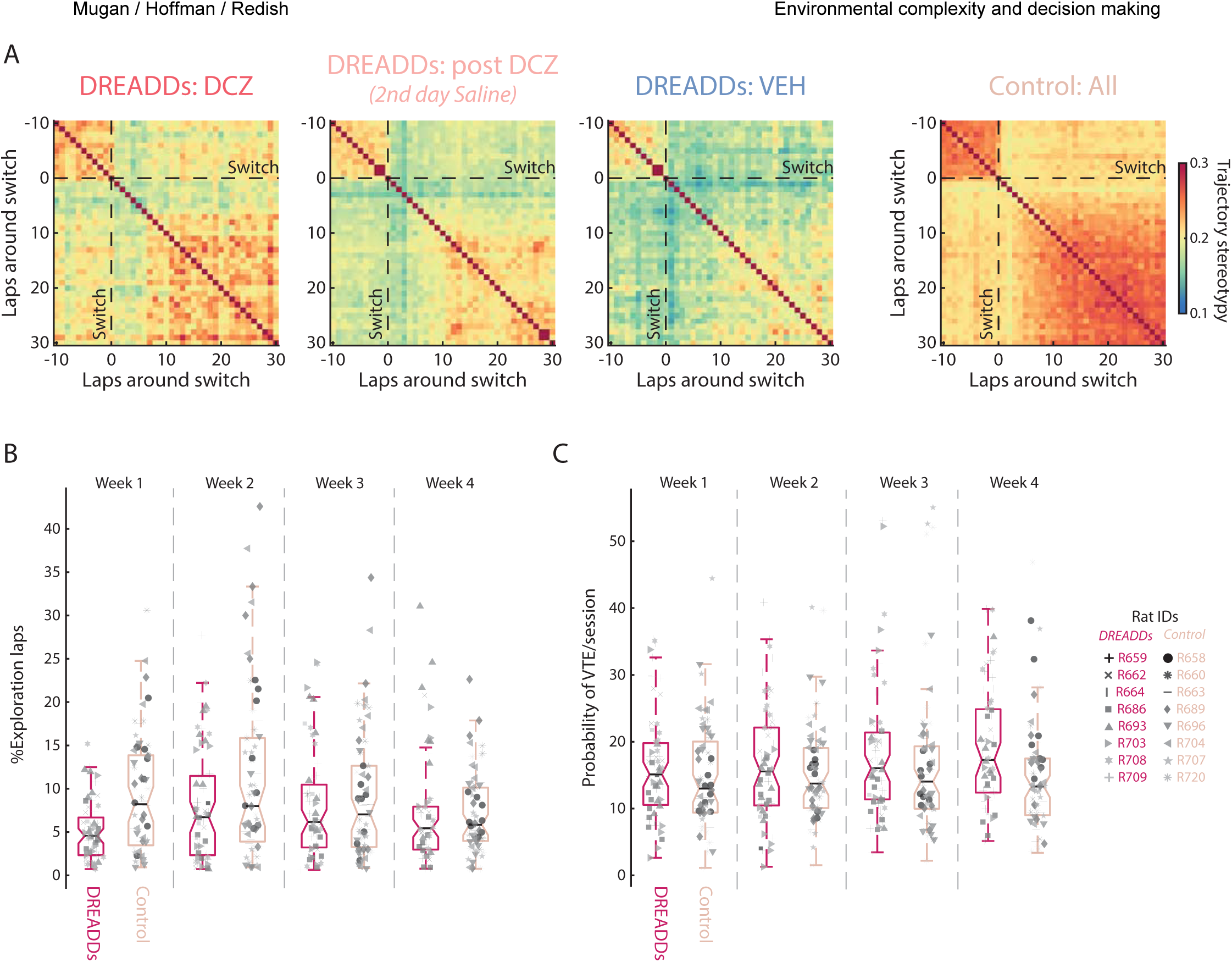
Long-term behavioral elfects of DREADD inactivation of dorsal mPFC, Related to Figure 8. **(A)** Same as Fig. 8J. Lap stereotypy aligned to task rule switches broken down by drug condition. Overall, DREADDs virus animals showed less stereotypy when compared to control virus animals. mPFC inactivation (DCZ) in DREADDs virus rats increased stereotypy; however, on the 2^nd^ day after both mPFC inactivation (post DCZ Saline and VEH, DREADDs rats displayed less behavioral stereotypy. **(B)** Similar representation to Fig. 8K. Distribution of the percentage of exploratory laps across the experimental weeks for DREADDs and control virus animals. Control virus animals changed their propensity to explore across weeks. **(B)** Similar to (B). Distribution of the proportion of VTE events across the experimental weeks for DREADDs and control virus animals.

**Supplementary Table 1:** Cell counts by rat and recorded region. Total, average, minimum, maximum, and standard deviation of the number of recorded units for each rat and brain region summed across all the sessions. Also showing the grand total of the number of recorded units for each brain region summed across all sessions and rats, as well as average, minimum, maximum, and standard deviation.

| **Rat ID** | **Total Sessions** | **Region** | **Total** | **Avg** | **Min** | **Max** | **Std** |
| --- | --- | --- | --- | --- | --- | --- | --- |
| R668 | 16 | *HC* | 588 | 36.75 | 20 | 53 | 9.38 |
|  |  | *mPFC* | 390 | 24.38 | 6 | 39 | 11.45 |
| R677 | 16 | *HC* | 1145 | 71.56 | 49 | 98 | 17.50 |
|  |  | *DLS* | 258 | 16.13 | 5 | 27 | 5.24 |
| R679 | 16 | *DLS* | 1037 | 64.81 | 37 | 90 | 14.87 |
|  |  | *mPFC* | 1111 | 69.43 | 48 | 93 | 15.86 |
| R699 | 16 | *HC* | 7 | 0.44 | 0 | 4 | 1.09 |
|  |  | *mPFC* | 38 | 2.38 | 0 | 7 | 2.16 |
| R710 | 16 | *HC* | 123 | 7.69 | 1 | 15 | 4.06 |
|  |  | *DLS* | 298 | 18.63 | 10 | 24 | 4.27 |
| R721 | 16 | *HC* | 999 | 62.44 | 36 | 93 | 16.41 |
|  |  | *mPFC* | 111 | 6.94 | 4 | 12 | 2.05 |
| R786 | 16 | *HC* | 1038 | 64.88 | 23 | 100 | 20.31 |
|  |  | *mPFC* | 583 | 36.44 | 31 | 43 | 3.97 |
| R787 | 16 | *HC* | 392 | 24.50 | 9 | 38 | 9.04 |
|  |  | *mPFC* | 254 | 15.88 | 8 | 31 | 7.49 |
| R788 | 16 | *DLS* | 700 | 43.75 | 32 | 59 | 8.20 |
|  |  | *mPFC* | 122 | 7.63 | 0 | 24 | 7.34 |

| **Regions** | **Total** | **Avg** | **Min** | **Max** | **Std** |
| --- | --- | --- | --- | --- | --- |
| ***HC*** | 4292 | 38.32 | 19.71 | 57.29 | 11.11 |
| ***DLS*** | 2293 | 35.83 | 21 | 50 | 8.15 |
| ***mPFC*** | 2609 | 23.30 | 13.86 | 35.57 | 7.19 |

